# Multivariate Time-Lagged Multidimensional Pattern Connectivity (mvTL-MDPC) for EEG/MEG Functional Connectivity Analysis

**DOI:** 10.1101/2024.01.20.576221

**Authors:** Setareh Rahimi, Rebecca L. Jackson, Olaf Hauk

## Abstract

Multidimensional connectivity methods are critical to reveal the full pattern of complex interactions between brain regions over time. However, to date only bivariate multidimensional methods are available for time-resolved EEG/MEG data, which may overestimate connectivity due to the confounding effects of spurious and indirect dependencies. Here, we introduce a novel functional connectivity method which is both multivariate and multidimensional, Multivariate Time-lagged Multidimensional Pattern Connectivity (mvTL-MDPC), to address this issue in time-resolved EEG/MEG applications. This novel method extends its bivariate counterpart TL-MDPC to estimate how well patterns in an ROI 1 at time point *t*_1_ can be linearly predicted from patterns of an ROI 2 at time point *t*_2_ while partialling out the multivariate contributions from other brain regions. We compared the performance of mvTL-MDPC and TL-MDPC on simulated data designed to test their ability to identify true direct connections, using the Euclidean distance to the ground truth to measure goodness-of-fit. These simulations demonstrate that mvTL-MDPC produces more reliable and accurate results than the bivariate method. We therefore applied this method to an existing EEG/MEG dataset contrasting words presented in more or less demanding semantic tasks, to identify the dynamic brain network underlying controlled semantic cognition. As expected, mvTL-MDPC was more selective than TL-MDPC, identifying fewer connections, likely due to a reduction in the detection of spurious or indirect connections. Dynamic connections were identified between bilateral anterior temporal lobes, posterior temporal cortex and inferior frontal gyrus, in line with recent neuroscientific models of semantic cognition.

## 1 Introduction

Cognitive functions depend on interactions among brain regions within distributed networks (Bullmore and Sporns, 2009; Passingham et al., 2002). Connectivity methods are essential to understand how areas interact across space and time to produce these processes. While functional magnetic resonance imaging (fMRI) is widely used for brain connectivity analyses, it only provides static pictures of brain networks, i.e. it has very limited temporal resolution (Glover, 2011). Therefore, functional connectivity methods suitable for EEG/MEG data are necessary to track cognitive processes non-invasively through time with millisecond temporal resolution. However, most widely used connectivity methods are limited in two ways, namely by assessing only bivariate and unidimensional connectivity. As in recent studies (Basti et al., 2019; Rahimi et al., 2023b, 2023a), we use the term “unidimensional” for methods that collapse signals to one time course per region, in contrast to “multidimensional” methods that use multiple time courses per ROI. We use the terms “bivariate” and “multivariate” to distinguish the computation of connectivity between only two regions at a time and when considering the effect of more than two regions simultaneously, respectively. Both of these factors determine the information gained from connectivity assessments. Unidimensional methods may lose important information within brain regions and therefore fail to identify true connections (Anzellotti et al., 2017b, 2017a; Basti et al., 2019, 2018) as interactions between brain areas are likely to be multidimensional (DiCarlo et al., 2012). Similarly, simplifying the complex interactions between different areas using multiple bivariate connectivity estimates is susceptible to mistaking indirect connections as direct ones. In other words, these bivariate measurements are prone to producing connections not as a result of a real, direct connection but because of their joint interaction with additional areas (Haufe et al., 2010; Iyer et al., 2020; Pozzi et al., 2021; Zalesky et al., 2012). As a result, functional connectivity networks may be overestimated (e.g. Iyer et al., 2020; Jackson et al., 2016; Rahimi et al., 2023a). Thus, to identify the key functional connections within a network without being over- or under-inclusive we require connectivity methods appropriate for high temporal resolution data that are both multivariate and multidimensional.

Multidimensional methods estimate relationships between time courses for multiple voxels per brain region (Anzellotti and Coutanche, 2018; Basti et al., 2020). For EEG/MEG applications, some multidimensional methods have been introduced operating in the frequency domain (Multivariate Interaction Measure (Ewald et al., 2012); Multivariate Lagged Coherence (Pascual-Marqui, 2007); and Multivariate Phase-Slope-Index (Basti et al., 2018)). However, while it is possible that some networks might only be reflected in specific frequency bands (Fries, 2015; Siegel et al., 2012), this may not be generalisable to all processes, such as early short-lived brain responses in event-related experimental paradigms. Additionally, most of these studies fail to provide detailed information about the dynamics of connectivity over time. Recently, Rahimi et al. (2023a) proposed Time-Lagged Multidimensional Pattern Connectivity (TL-MDPC) which provides a multidimensional method for event-related responses in EEG/MEG data in the time domain. This bivariate undirected functional connectivity method estimates how well patterns of responses at a specific time point in one ROI can predict those of another ROI at another time point using cross-validated regularised ridge regression (Rahimi et al., 2023a). As a result, the method estimates connectivity between pairs of brain regions over time and across different time lags. TL-MDPC captured connectivity missed by unidimensional methods in both simulated and real EEG/MEG data (Rahimi et al., 2023a). However, this method is currently bivariate only.

Multivariate methods address the overly inclusive nature of bivariate methods by including all relevant regions in one larger connectivity model, rather than estimating a separate bivariate model for each pair of regions (Haufe et al., 2010). A number of multivariate effective (e.g. Baccalá and Sameshima, 2001; Barnett and Seth, 2014; Blinowska et al., 2004; Franaszczuk et al., 1985; Kus et al., 2004; Talebi et al., 2019) and functional (e.g. Salvador et al., 2020; Valdés-Sosa et al., 2005) connectivity methods have been proposed, yet they are all unidimensional and reduce the activity in each region to a single value per time point. No multivariate multidimensional connectivity method has been introduced yet. Here, we extend TL-MDPC (Rahimi et al., 2023a) to assess multivariate as well as multidimensional connectivity in EEG/MEG data. This means we take into account all relevant ROIs when computing the connectivity between patterns of responses in pairs of ROIs. We first estimate a coefficient matrix, mapping the vertices of a target ROI (ROI *X*_1_) at time point *t*_1_ to all other ROIs at time point *t*_2_, then choose the relevant subset of coefficients for a corresponding pair of ROIs (here ROI *X*_2_ to ROI *X*_1_), and finally estimate how well patterns in ROI *X*_2_ can predict patterns in ROI *X*_1_ in terms of their partial explained variance. While multidimensional connectivity methods have been shown to capture more information than unidimensional ones, a multivariate connectivity method is expected to show fewer connections than its bivariate counterpart (both fewer pairs of regions and connectivity at fewer time points), because of its insensitivity to indirect and spurious connections. Thus, a multivariate method is likely to produce fewer connections than a bivariate approach, but those connections are more likely to be true direct. Thus, the output can be expected to be more reliable and meaningful.

We evaluated the method’s performance using simulations and a real EEG/MEG dataset. We used simulated data with a range of signal-to-noise ratios (SNRs) to find out how sensitive mvTL-MDPC is to 1) false positive errors for random noise, 2) one-to-one bivariate dependencies, 3) multivariate dependencies with different transformation strengths, 4) all-to-all multivariate dependencies, 5) indirect connections, and 6) spurious connections. In the following, ‘spurious connections’ refer to the case where one region affects two others, potentially creating a false connection between the latter. This is distinct from ‘indirect connections,’ where a chain of influence can be mistaken for a direct link. We also applied this new method to an EEG/MEG dataset, contrasting two tasks requiring different depths of semantic processing of a single word, and compared the resulting connectivity between regions of the semantic network (left and right anterior temporal lobes (ATL), posterior temporal cortex (PTC), inferior frontal gyrus (IFG), angular gyrus (AG), and primary visual area (PVA)) to the bivariate results from our previous study (Rahimi et al., 2023a).

## 2 Material and Methods

We here extend our previous bivariate TL-MDPC method to the multivariate case, allowing it to consider several regions’ patterns simultaneously within one model, aiming to better distinguish direct from indirect or spurious connections between regions. Targeted simulations and real EEG/MEG data were utilised to compare this new method to the bivariate approach. We hypothesised that the multivariate method would be more robust against indirect and spurious connections whilst still identifying pairs of regions which directly interact. We first present the shared logic and procedures between the two methods. For both methods, we 1) prepare pattern of responses at each time point in every region, 2) find a model that transforms patterns from one region at one time point to patterns in another region at another time point, 3) predict the target ROI patterns, 4) and quantify the connectivity measure by computing the explained variance (EV) between the real and predicted output patterns.

Let us consider ***X*_1_** and ***X*_2_** to be the brain activity patterns at time points 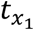 and 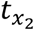, which are of size 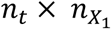 and 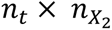, respectively, where *n*_*t*_ is the number of trials, and 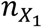 and 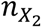 are the number of vertices in the two regions. We want to figure out if there exists a vertex-to-vertex transformation between activity patterns at different time points. In other words, we intend to estimate how well ***X***_**2**_ at time point 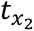 can predict ***X***_**1**_ at time point 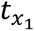 through a linear transformation.

To prepare the patterns and build a model, we first need to deal with the issue of spatial resolution. EEG/MEG signals are inherently blurry and have limited spatial resolution (Hauk et al., 2019; Palva et al., 2018). This leads to correlated signals across vertices within and across ROIs. Thus, different vertices carry redundant information. To address this issue while preserving the genuine space of activation, we previously proposed a “feature selection” solution using unsupervised k-means clustering to sub-sample the most informative vertices within each ROI (Rahimi et al., 2023a). Every pattern matrix consists of brain activity across trials and across vertices. Here, vertices serve as samples or observations and trials as features. To do the subsampling, we cluster all vertices and then pick the vertex with the highest variance as the most informative one. Clustered patterns of ***X*_1_** and ***X*_2_** are now of size 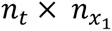 and 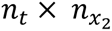, where 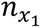 and 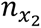 are the number of clusters in ROI *X*_1_ and ROI *X*_2_.

Having prepared the dimensionality-reduced pattern matrices, we can estimate the transformation between patterns at different time points, using cross-validated regularised ridge regression (Hoerl and Kennard, 1970). After extracting a transformation ***T*** for two specific patterns, we can predict patterns in the target ROI *X*_1_ (***X̂***_**1** ***test***_) through ***T*** and test subset ***X***_**2** ***test***_. We then quantify the connectivity by computing the explained variance between the real output, ***X***_**1** ***test***_, and the predicted output, ***X̂***_**1** ***test***_ Note that for the bivariate method, we follow Rahimi et al., (2023a) by performing this procedure in both directions, as both are very similar and to avoid any potential bias due to there being different numbers of vertices and different amounts of noise, i.e. ***X***_**1**_ being predicted from ***X***_**2**_ and ***X***_**2**_ being predicted from ***X***_**1**_, and report the average of resulting EVs. However, for a fairer comparison with the multivariate method (where results differed more between the two directions), we also check the effect of only including predictions of later from earlier time points. The EV metric can vary from 1, suggesting a strong linear dependency, to zero (and even negative values) suggesting no dependency at all. Since both the negative values and zero are an indication of no dependency between patterns we replace negative values with zero.

For every pair of ROIs, these steps are performed for all pairs of time points, and the resulting EVs are placed in a time-time matrix referred to as a Temporal Transformation Matrix (TTM; Rahimi et al., 2023a). This provides the time points of ROI *X*_2_ in the vertical axis and that of ROI *X*_1_ in the horizontal axis. In every TTM, the diagonal entries show the simultaneous connectivity, while other parts reflect the lagged connectivity between the two ROIs, with the upper diagonal representing connectivity where ***X***_**1**_ is ahead in time, and the lower diagonal representing connectivity where ***X***_**2**_ is ahead.

### 2.1 The mvTL-MDPC method

Both methods compute the relationship between two areas, yet they differ with respect to the consideration of information in additional ROIs. For the bivariate method, to estimate the connectivity between ROI X_1_ and ROI X_2_, we only consider the patterns of these two regions, as shown in Figure 1a. In contrast, for the multivariate method we consider the patterns of all selected regions when determining this relationship, as shown in Figure 1b.

**Figure 1.**
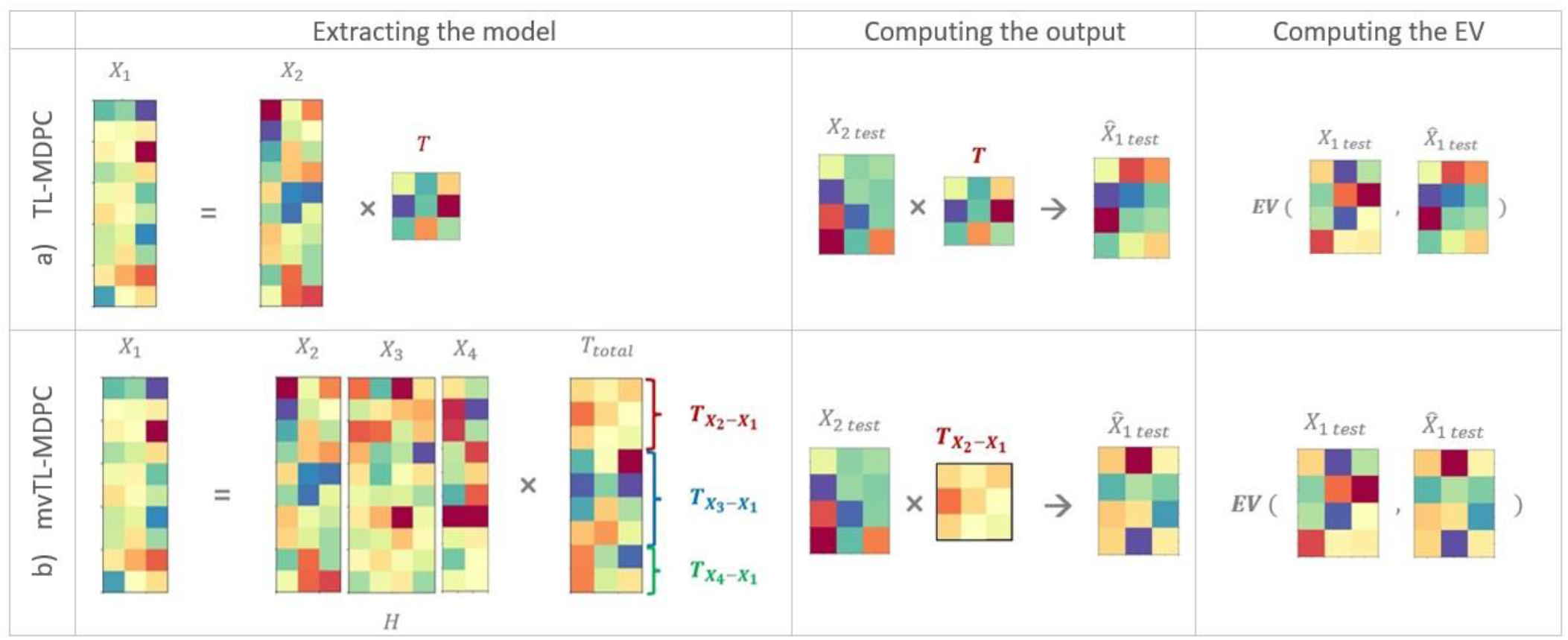
Illustration of bivariate and multivariate multidimensional pattern connectivity (MDPC) approaches. A) The principle of the bivariate MDPC method is displayed (as in Rahimi et al., 2023a). We assess the relationship between activity patterns in ROI X_2_ and ROI X_1_ at two different time points (here, only a single time lag is shown), using ridge regression. Each matrix indicates activity patterns in one ROI at one time point, with rows in each matrix indicating activation across different trials, and columns representing activation over different vertices in the ROI. B) Illustration of mvMDPC and how it detects the linear transformations ***T*** between patterns of ROI X_2_, X_3_, and X_4_ and ROI X_1_, using ridge regression (this is an example for a network of four ROIs). To understand the relationship between ROI X_2_ and ROI X_1_ the mapping with all regions is estimated within this single model and then the relevant portion of the mapping is considered, thus, the portion of variance that is better explained by relationships with other regions, is removed. This was hypothesised to reduce the likelihood of finding no relationship in cases where there is no true direct connection, such as when there are indirect or spurious connections, while still identifying true direct connections. For both approaches, we report the resulting explained variance (EV) between the predicted and real patterns of the target ROI (ROI X_1_).

Let us consider that ***X***_**1**_, ***X***_**2**_, …, and ***X***_***n***_ are dimensionality-reduced patterns of responses within ROI X_1_, ROI X_2_, …, and ROI X_*n*_ at time point *t*_1_, for ***X***_**1**_, and *t*_2_ for the rest of the patterns, and of size *n*_*t*_ × *n*_*x*1_, *n*_*t*_ × *n*_*x*2_, …, and *n*_*t*_ × *n*_*xn*_, respectively, with *n*_*t*_ being the number of trials, and *n*_*xi*_ the number of vertices in ROI X_*i*_. We want to find out whether there is a mapping between the two patterns ***X*_1_** and ***X*_2_** at two time points *t*_1_ and *t*_2_, but we want to consider that this relationship may also be affected by connections with regions X_3_ to X_*n*_. To assess this, we need to build a model to extract the mapping between the two patterns of interest in the presence of the other areas. mvTL-MDPC creates this model by estimating the effect of all other patterns on ***X***_**1**_, each potentially explaining a portion of the target pattern’s variance, and then extracting the contribution of ***X***_**2**_. A general mapping between ***X***_**1**_ and the rest of the patterns can be described as follows:

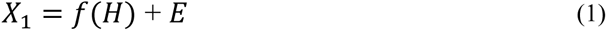

Where *H* is [***X***_**2**_, ***X***_**3**_, …, ***X***_***n***_] and of size *n*_*t*_ × *n*_*h*_, *n*_*h*_ = *n*_*x*2_ + ⋯ + *n*_*xn*_. The linear transformation, i.e., the mapping of the patterns in all the ROIs into ***X***_**1**_, can be written as:

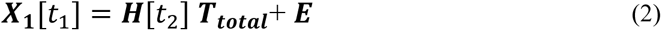

Then, the total transformation ***T***_***total***_ can be estimated using regularised Ridge Regression (Hoerl and Kennard, 1970) and the train subset of trials:

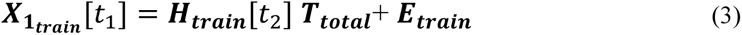

where ***T***_***total***_ is the total transformation matrix, of size *n*_*h*_ × *n*_*x*1_ and ***E***_***train***_ is the error matrix of size *n*_*t*_ × *n*_*x*1_. ***T***_***total***_ can be estimated using the regularised pseudoinverse of ***H***_***train***_:

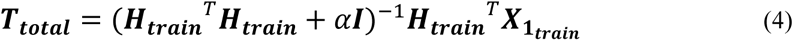

where *⍺* is the regularisation parameter (Tikhonov and Arsenin, 1977) to be determined using cross-validation and ***I*** is the identity matrix of size *n*_*h*_ × *n*_*h*_.

Once the ***T***_***total***_ is extracted, we can estimate the connectivity between ROI X_1_ and any other ROI, e.g., ROI X_2_, using their own relevant transformation sub-matrix, namely 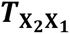, being a subset of ***T***_***total***_. Afterwards, we can predict the patterns in ROI X_1_ using only the portion that is the result of the transformation of activation from ROI X_2_ on this region, using the test subset and 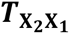:

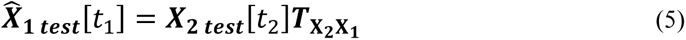

where ***X̂***_**1** ***test***_ is the predicted pattern. For each vertex j=*1*, …, *n*_*x*1_, we then compute the explained variance (EV):

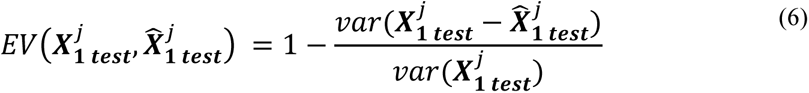

Finally, as a metric for the multidimensional connectivity strength between two ROIs, we average the EVs across vertices:

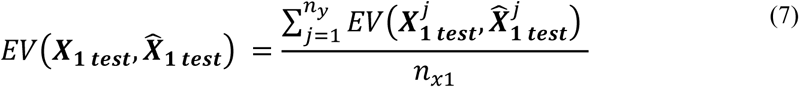

The highest possible score of 1 suggests strong connectivity while values close to or below zero indicate weak connectivity. As with the bivariate method any negative value is replaced by zero: *max* (*EV*(***X***_**1** ***test***_, ***X̂***_**1** ***test***_), 0). For the bivariate method, the resulting EVs were similar in both directions and therefore the prediction is performed in both directions and combined as described above to maintain consistency with Rahimi et al., (2023a). However, in the multivariate case the models specified in equation (2) (matrices ***X*** and ***H***) differ markedly between the two approaches, therefore here we only predict future patterns from past patterns. To ensure this is not the cause of any differences between the methods we ran additional analyses where the bivariate case is only applied in this direction. We found that the effect of direction did not lead to meaningful differences and could not explain the critical differences between the results of the two methods (see Results).

### 2.2 Simulations: Performance of bivariate and multivariate TL-MDPC for different relationships between patterns

We assessed mvTL-MDPC’s ability to capture ground truth connectivity in comparison to TL-MDPC’s performance, initially in specially-designed simulated connectivity scenarios and then across thousands of random networks, considering an optimal range of trials, vertices, and SNRs. This section begins with an evaluation of these methods in targeted scenarios to clarify our conclusions, followed by a broader analysis in random networks to enhance the generalisability of our findings. As we are not simulating time courses here but simply estimating pattern relationships across trials per sample latency, we use the shorter terms ‘MDPC’ and ‘mvMDPC’ for this section.

We contrast MDPC and mvMDPC on simulated data for six different scenarios (see Figure 2):

1. no relationship between the regions,
2. a bivariate linear multidimensional relationship between two regions,
3. multivariate multidimensional relationships with different strengths (multiple regions affecting one region, each with a different contribution),
4. multivariate linear multidimensional relationships with an all-to-all mapping within a network,
5. ‘indirect’ connections, and
6. ‘spurious’ connections.

**Figure 2.**
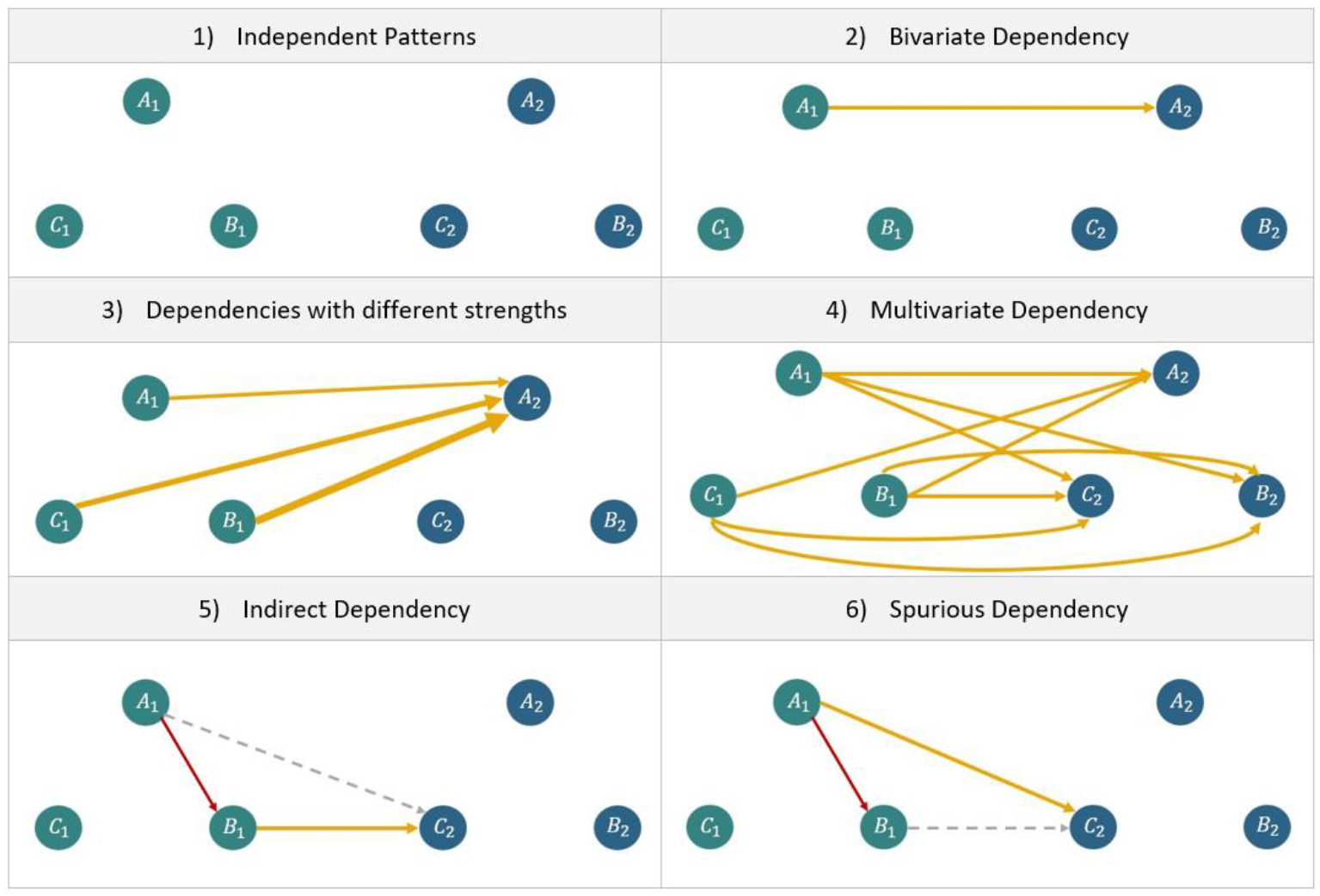
Representation of the different simulation scenarios. The regions *A, B,* and *C* contain patterns of responses at time point *t*_1_ and *t*_2_, respectively. 1) All patterns are independent with no reliable transformation between the regions and as a result no connectivity. 2) There is one bivariate (one-to-one) multidimensional relationship between *A*_1_ and *A*_2_ through a linear transformation ***T***. 3) There are relationships between all regions at time point *t*_1_ and one region at time point *t*_2_, each with a different transformation strength. 4) There exist multivariate (all-to-all) relationships between all regions at two time points. 5) There is one zero-lag dependency, *A*_1_-*B*_1_, as well as a lagged dependency, *B*_1_-*C*_2_, creating an ‘indirect connection’ where *A*_1_-*C*_2_ may be identified as connected although A does not directly influence C. 6) There is one zero-lag dependency, *A*_1_-*B*_1_, as well as a lagged dependency, *A*_1_-*C*_2_, which may lead to the identification of a ‘spurious connection’, *B*_1_-*C*_2_. Although *B* never influences *C* or vice versa there is a relationship between their activity patterns due to the shared influence of a third region. Note that indirect and spurious connections may occur across time, across regions, or both and do not specifically require one lagged and one zero-lagged connection. Direct lagged connections are yellow, zero-lag relationships (relationships between ROIs at the same time point) are red, and indirect or spurious connections that do not reflect true direct connections are dashed grey lines.

Here we use the term ‘spurious connections’ specifically when region A influences both regions B and C, and a dependency between B and C may be identified without a true connection. This allows us to distinguish this scenario from ‘indirect connections’ where region A influences region B, and region B influences region C without A and C being directly related, where the indirect dependency between A and C may be misidentified as a direct connection.

Although described across regions, these scenarios can also be interpreted as relating to the same areas across different time points. Thus, both indirect and spurious connections could occur between areas that are not directly connected or between areas that are directly connected but not at the time point assessed. Both would result in an overly inclusive network and misrepresent the network dynamics. For each scenario, we consider three regions at an arbitrary time point *t*_1_ (*A*_1_, *B*_1_, *C*_1_), and the same regions at another arbitrary time point *t*_2_(*A*_2_, *B*_2_, *C*_2_). For each set of parameters (e.g., different levels of SNR), we computed the functional connectivity with MDPC and mvMDPC and compared each result with the true connectivity matrix (the ground truth) using Euclidian distance as the goodness-of-fit to contrast the performance of the two methods. The first two scenarios are simple checks for cases when there is no connection or a simple bivariate connection, and we would expect both methods to do well. The second case also helps determine whether the multivariate method is simply worse at identifying connections than the bivariate method, in which case it would perform worse. In cases 3 and 4 we consider different kinds of multivariate relationships, where the multivariate method is expected to identify a solution closer to the ground truth. The final two cases include the types of spurious and indirect relationships which are typically considered likely to be highlighted as strong connections using bivariate methods, making them indistinguishable from the critical direct connections. We hypothesise the multivariate method will be less sensitive to these issues. After assessing these specific simulation patterns, we compare our methods using networks with randomly generated connections (i.e., networks without an imposed structure) to determine whether any identified benefits of the multivariate method are generalisable across a range of network architectures. In scenarios where there are dependencies, different levels of noise were introduced by varying the standard deviation in a zero-mean normal distribution.

#### 2.2.1 Scenario 1: Checking for false positive connections between two independent activity patterns

Figure 2-1 shows the first scenario, indicating two independent sets of patterns with no connectivity, i.e., 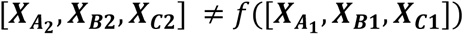. Thus, the EV should be close to zero for all possible pairs. To create these patterns, six independent matrices were generated using normal distributions (*mean*=0 and *std*=1).

#### 2.2.2 Scenario 2: Testing the methods’ ability to capture connectivity between two regions with a bivariate dependency between their patterns

Figure 2-2 represents scenario 2, where there are simple one-to-one mappings between the activity patterns in one region at the first time point and another at the second, reflecting a direct, bivariate linear multidimensional relationship. The vertices in *A*_1_ are uncorrelated to each other yet transformed to *B*_2_ through a matrix ***T***, so that 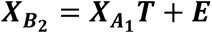, where ***T*** is of size 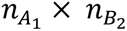. We generated five independent patterns 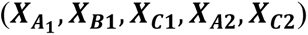 using normal distributions (*mean*=0, and *std*=1). For transformation matrix ***T***, mapping 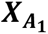 to 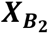, we used a normal distribution (*mean*=0, and *std*=1). 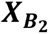 was then computed through multiplying 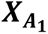 and ***T***. The EV should be high for the connected pair and close to zero for all others.

#### 2.2.3 Scenario 3: Testing the sensitivity of each method to multivariate dependencies with different strengths

Figure 2-3 illustrates scenario 3 in which one region’s patterns are created from multiple other regions’ patterns through transformation matrices with different overall strengths, so that: 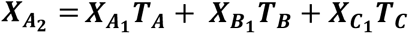, where mean(***T***_***C***_)> mean(***T***)> mean(***T***_***A***_). Here we generated five independent patterns (reflecting all the regions at the first time point and two unconnected regions at the second; 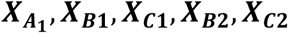) using normal distributions (*mean*=0, and *std*=1), as well as three transformation matrices, mapping 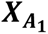 to 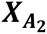 (*mean*=0, and *std*=1), ***X***_***B*****1**_ to 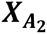 (*mean*=1, and *std*=1), and ***X***_***C*****1**_ to 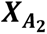 (*mean*=2, and *std*=1). The pattern in the final area at the second time point, 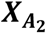 is then computed through multiplying 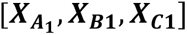 and the three coefficient matrices. The EVs should reflect the relationship between the one connected area at the second time point and each area at the first time point. They should also accurately reflect the differences in strength between these three identified connections.

#### 2.2.4 Scenario 4: Testing the ability of each method to capture multivariate all-to-all dependencies between patterns

Figure 2-4 shows the multivariate multidimensional relationships in scenario 4 where all regions within the network are connected to each other, so that 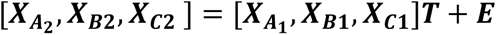, where ***T*** is of size 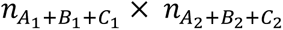. We generated three independent patterns 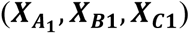 using normal distributions (*mean*=0, and *std*=1), as well as a coefficient matrix ***T*** based on a normal distribution (*mean*=0, and *std*=1). As a result of applying this transformation, each of the patterns at the first time point contributes to the pattern in all the regions at the second time point. The EVs should be high for all pairs of regions assessed across time.

#### 2.2.5 Scenario 5: Testing the likelihood of each method to falsely identify indirect dependencies

Figure 2-5 indicates scenario 5 where there is an indirect relationship between a pair of regions. In this scenario, there is one zero-lag dependency between *A*_1_ and *B*_1_, as well as a lagged dependency (i.e., a relationship between the patterns at two different time points as in the prior scenarios) between *B*_1_ and *C*_2_. A zero-lag dependency refers to simultaneous dependencies between two regions, which could reflect leakage or a relationship between the current patterns could result from true connectivity (or indirect or spurious connectivity) at an earlier time. Accordingly, there is an indirect dependency between *A*_1_ and *C*_2_, as there is no direct connection between these areas, yet a relationship may be identified due to the influence of region *A* on region *B*, combined with the influence of region *B* on region *C*. Equivalent patterns are also possible across different regions within a single time point or within the same pair of regions across different time points. To assess this, four independent patterns 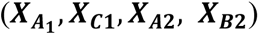 were generated using normal distributions (*mean*=0, and *std*=1), as well as two coefficient matrices, mapping 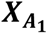 to ***X***_***B*****1**_ (zero-lag dependency), and ***X***_***B*****1**_ to ***X***_***C*****2**_ (lagged dependency), through a normal distribution (*mean*=0, and *std*=1). In this case *B*_1_-*C*_2_ is a direct connection, while *A*_1_-*C*_2_ is an indirect connection. Thus, ideally the EVs would reflect the true situation with strong values between *A* and *B,* and *B* and *C* and lower values for the connection between *A* and *C*. However, the distinction between direct and indirect connections is hard to detect and bivariate methods, in particular, are expected to produce high EVs for both.

#### 2.2.6 Scenario 6: Testing the likelihood of each method to falsely identify spurious dependencies

Figure 2-6 shows a spurious connection. In scenario 6, the existence of a zero-lag connection between *A*_1_ and *B*_1_(caused either by leakage or prior interaction of these regions), and a true lagged connection between *A*_1_ and *C*_2_, could lead to the identification of a spurious connection between *B*_1_ and *C*_2_. Here regions *B* and *C* are not connected, yet both are influenced by the activity in region *A* leading to some relationship between their patterns. This differs from scenario 5 as here the activity in *B* has no influence on the pattern in *C* even indirectly, yet this connection may still be identified. As with indirect connections, these spurious connections can occur between regions that are never directly connected or regions that are connected but not at that time. To assess this we generated four independent patterns 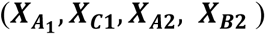 using normal distributions (*mean*=0, and *std*=1), as well as two coefficient matrices, mapping 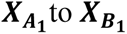 to 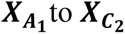, through a normal distribution (*mean*=0, and *std*=1). The EVs should be high for the direct connections between *A* and *B* and *A* and *C*, but low for *B* to *C*. However, this distinction is expected to be hard to detect, particularly for the bivariate method.

#### 2.2.7 Simulation Parameters

To evaluate the performance of mvMDPC versus the bivariate method, we computed the connectivity in each scenario over realistic ranges of SNRs for 200 trials and 10 vertices within each ROI. We used 10 vertices as a reasonable number within the range of minimum (5) and maximum (13) number of vertices obtained from the previous application of our feature selection approach on the EEG/MEG dataset used here (Rahimi et al., 2023a). Different levels of noise were introduced from a zero-mean normal distribution with a varying standard deviation (*std*=10*^std_pow^*, where *std_pow* ∈[−2, −1.5, −1, −0.5, 0, 0.5, 1, 1.5, 2]). For every set of parameters, we ran 500 simulations and reported the average as the final connectivity value. At each simulation, we computed the Euclidian distance between the (3 by 3) ground truth matrix and the (3 by 3) MDPC result, as well as the ground truth and the (3 by 3) mvMDPC result to contrast these distances using a paired t-test.

#### 2.2.8 Random Network Simulations

The previous simulations provided insight into each methods’ behavior when applied to specific connectivity scenarios. Here, we provide a more exhaustive comparison of the methods extending beyond these particular scenarios, by comparing the two methods over a wider range of connectivity patterns using a large number of random networks. This improves the generalisability of the findings to a broader set of network types. These networks consisted of three regions at two time points as above, yet their connectivity varied as the following parameters were selected randomly: 1) the mean of the normal distribution used to generate patterns and transformation matrices, in the range of [−2.5, 2.5], 2) the standard deviation of the normal distribution used to generate the patterns and transformation matrices, in the range of [1,6], 3) the number of vertices (an integer between 5 and 15), 4) the number of trials (an integer between 110 and 250), 5) the number of connections (each connection has a 0.5 likelihood of being present), and 6) the sparsity of the transformation matrices (i.e., the percentage of non-zero elements in the matrix, a number between 0 and 100%). We replicated this procedure 10000 times. For each replication we computed the mvMDPC and MDPC results and their Euclidian distances from the ground truth. We performed two sets of these simulations, with the second set including zero-lag dependencies as well as connections between regions over time. We considered all 6 possible zero-lag dependencies (e.g., from *A*_1_ to *B*_1_: *B*_1_ = *B*_1*prime*_ + *A*_1_*T*_*A*1*B*1_ where *B*_1*prime*_ and *A*_1_ are independent) with the same likelihood (0.5)). As before these zero-lag dependencies may be artefactual, reflecting the limited resolution of EEG/MEG data or they may be the consequence of the prior interaction of the regions.

### 2.3 Comparing bivariate and multivariate TL-MDPC in a real EEG/MEG dataset

To compare the two methods on real data, we also applied mvTL-MDPC to an existing EEG/MEG dataset, contrasting the brain connectivity of two tasks with different semantic processing demands: a semantic decision task (SD), requiring the retrieval of detailed semantic features of a word’s meaning, and a lexical decision (LD) task, requiring only the distinction between existing and non-existing words and therefore shallower semantic processing. We can then compare our new results with those from a previous study (Rahimi et al., 2023a).

#### 2.3.1 Participants

We employed the same dataset as in our prior studies (Rahimi, 2023; Rahimi et al., 2023a, 2022) containing EEG/MEG recordings of 18 healthy native English speakers (mean age 27.00±5.13, 12 female) with normal or corrected-to-normal vision (for more detail see Farahibozorg, 2018; Rahimi et al., 2022). The experiment was approved by the Cambridge Psychology Research Ethics Committee and volunteers were paid for their time and effort.

#### 2.3.2 Stimuli and Procedure

The analysis was applied to EEG/MEG data for 250 words presented in four randomly presented blocks which also contained 250 pseudowords. Three blocks included a semantic decision task (SD), and one a lexical decision task (LD). In the LD task, participants were required to identify whether the presented stimulus was referring to a word or a pseudoword, while in the SD task, they needed to decide whether the presented stimuli were a member of a particular semantic category (targets), specifically “non-citrus fruits”, “something edible with a distinctive odour” and “food containing milk, flour or egg”. The stimulus set for SD included 30 target words that required a button press response. As in previous analyses of this dataset, we only analysed the non-target words. The duration of stimulus presentation and average SOA were 150ms and 2400ms, respectively.

#### 2.3.3 Data Acquisition and Pre-processing

Data acquisition and pre-processing steps are identical to previous analyses of this dataset (Farahibozorg, 2018; Rahimi et al., 2023a, 2022). MEG and EEG signals were recorded simultaneously using a Neuromag Vectorview system (Elekta AB, Stockholm, Sweden) and MEG-compatible EEG cap (EasyCap GmbH, Herrsching, Germany) at the MRC Cognition and Brain Sciences Unit, University of Cambridge, UK. MEG was recorded using a 306-channel system including 204 planar gradiometers and 102 magnetometers. EEG was acquired by a 70-electrode system with an extended 10-10% electrode layout. Data were sampled at 1000Hz.

MEGIN Maxwell-Filter software was used to implement signal source separation (SSS) to remove distant sources of noise and to alleviate small head movement effects (Taulu and Kajola, 2005). The MNE-Python software package (Gramfort et al., 2014, 2013) was used to perform the pre-processing and source reconstruction steps. For each participant, the raw data was visually inspected and bad EEG channels were marked for interpolation. We then applied a finite-impulse-response (FIR) bandpass filter (0.1 and 45 Hz) and FastICA algorithm to remove eye movement artefacts (Hyvarinen, 1999; Hyvärinen and Oja, 2000). Epochs were obtained using time intervals from 300ms pre-stimulus to 600ms post-stimulus.

#### 2.3.4 Source Estimation

To reconstruct the source signals, we employed L2-Minimum Norm Estimation (MNE) (Hämäläinen and Ilmoniemi, 1994; Hauk, 2004). To do so, the inverse operators were assembled using a 3-layer Boundary Element Model (BEM) of the head geometry yielded from structural MRI images, with the assumption of sources being perpendicular to the cortical surface (a “fixed” orientation constraint). Noise covariance matrices were constructed using 300ms-baseline periods, after identifying the best of the different methods included in MNE python (’shrunk’, ‘diagonal_fixed’, ‘empirical’, ‘factor_analysis’) (Engemann and Gramfort, 2015). We regularised the inverse operator for evoked responses using MNE-Python’s default SNR = 3.0. Finally, the source estimates of each participant were morphed to the standard Freesurfer brain (fsaverage).

#### 2.3.5 Regions of Interest

As in our previous study we used the Human Connectome Project (HCP) parcellation (Glasser et al., 2016) to select six regions of interests (ROIs) associated with semantic processing in the visual domain based on the literature, namely left ATL, right ATL, left PTC, left IFG, left AG and left PVA.

#### 2.3.6 Leakage

EEG/MEG source signals can suffer from leakage and as a result provide limited spatial resolution, which can confound connectivity analyses. As a result, here our analyses are interpreted with reference to the leakage matrix previously constructed for this dataset (Rahimi et al., 2023a, 2022). In these studies a leakage matrix was created to describe the amount of possible leakage between our six ROIs. First, the resolution matrix was computed from which the desired point spread and cross-talk functions (PSFs and CTFs, which are the same for MNE) (Hauk et al., 2011; Liu et al., 2002) for all regions can be extracted. By multiplying the PSF/CTFs and non-homogeneous activation vectors, the leakage from one region to another is obtained. This procedure was repeated 100 times for each PSF and participant, and the average results were used in the next step. As shown in Figure 3a, the leakage activation is summarised by taking the absolute values and summing them across all vertices.

**Figure 3.**
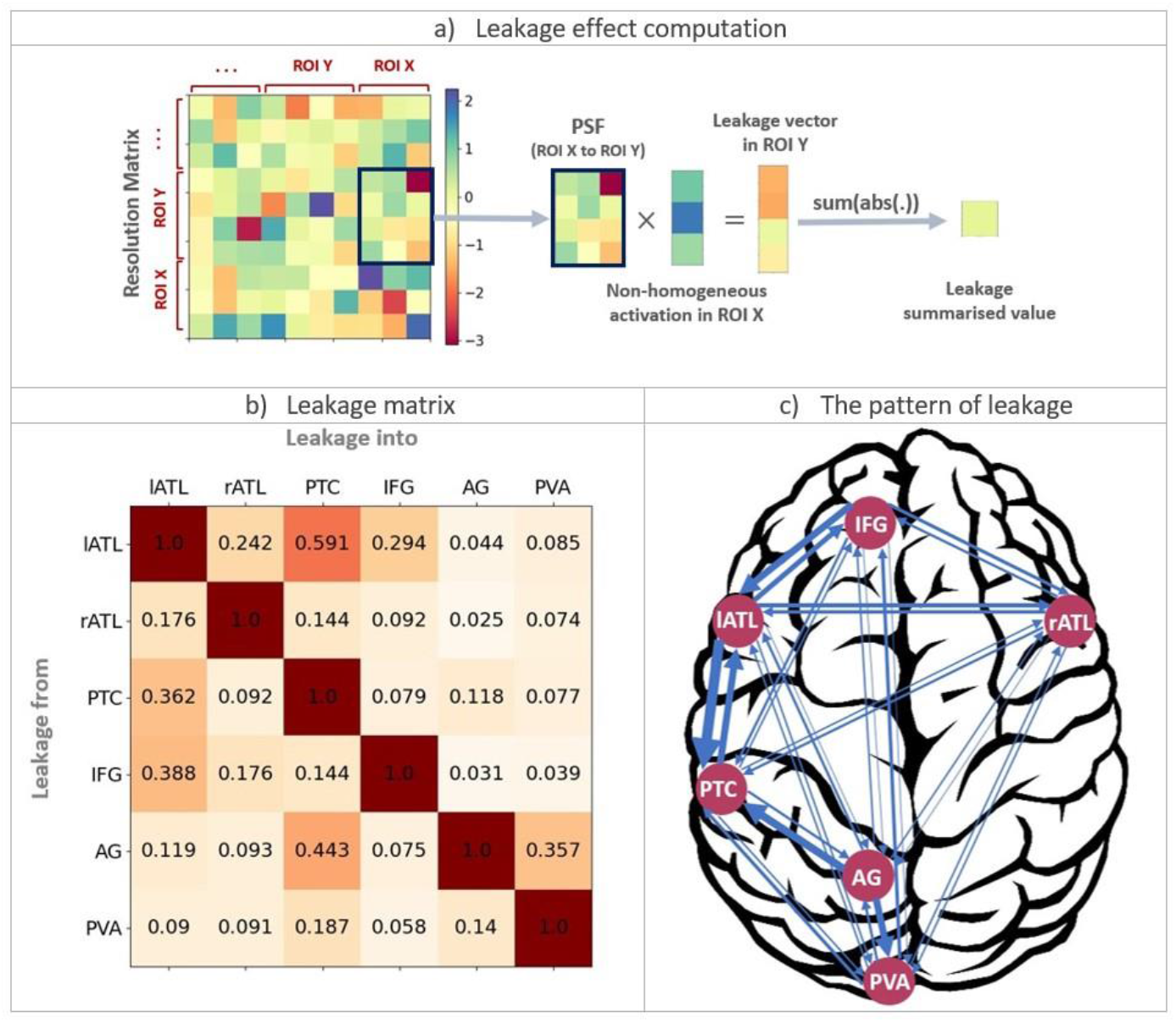
Quantitative source leakage evaluation of the real EEG/MEG dataset. a) Leakage indices based on point spread/ cross-talk functions (PSF/CTFs) for each pair of regions. b) Leakage matrix, indicating leakage indices between each pair of regions in the EEG/MEG analysis. c) The pattern of leakage across the regions used in our analysis. The width of the arrows reflects the leakage indices in b) (obtained from Rahimi et al., 2023a).

Figure 3b shows the leakage matrix containing leakage indices for pairs of ROIs, defined as the leakage from the origin ROI to the target ROI divided by the leakage from the target ROI to itself. To provide a clear interpretation of the matrix, the leakage values between 0-0.2/0.2–0.4/0.4–0.6/0.6–0.8/0.8-1 are labelled as low/low-medium/medium/medium-high/high. The leakage matrix suggests that all values, except for the leakage from lATL to PTC, represent medium or low leakage.

#### 2.3.7 Application of mvTL-MDPC to real EEG/MEG data

We calculated the mvTL-MDPC for every pair of ROIs at every 25ms from 100ms pre-stimulus to 500ms post-stimulus, to predict current patterns of ***X***_**1**_ from past patterns of ***X***_**2**_, and reported the resulting EV as the connectivity metric. We then presented all EVs at different lags for each pair, in a TTM (Rahimi et al., 2023a). In any TTM, the upper diagonal elements show how well the pattern of the ROI on the horizontal axis predicts the pattern of the ROI on the vertical axis, and the lower diagonal represents how well the pattern of the ROI on the vertical axis predicts the pattern of the ROI on the horizontal axis.

#### 2.3.8 Statistical analysis

As the SD task has three different blocks and the LD task has one, the TTMs of each SD block were computed separately and then averaged, to have a fair and unbiased comparison with the LD TTMs (Bastos and Schoffelen, 2016). The SD and LD TTMs were then compared using cluster-based permutation tests, implemented in MNE python (Maris and Oostenveld, 2007). A two-tailed t-test was applied with an alpha-level of 0.05 and 5000 randomised repetitions. To lessen the effect of spurious or small clusters, only clusters with size greater than 2% of the TTMs total size (12) were reported.

## 3 Results

### 3.1 Simulation results

#### 3.1.1 Scenario 1: Testing for false positive connections between two independent activity patterns

Figure 4 indicates the connectivity scores for each pair of regions based on the ground truth, mvMDPC and (bivariate) MDPC, averaged across all simulation runs, for the case where all patterns are unrelated (as in Figure 2-1). For both methods, all values are close to zero, suggesting that neither method is prone to false positive errors when no relationship exists.

**Figure 4.**
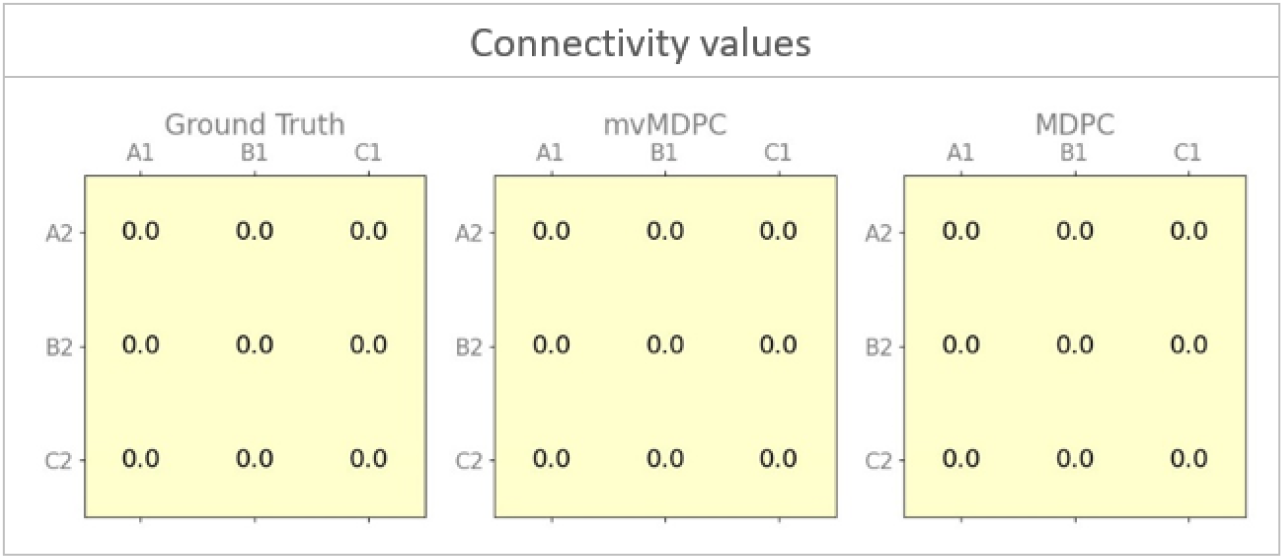
Simulation results for independent patterns. It shows connectivity values (explained variance) for the ground truth, mvMDPC, and MDPC methods. Both methods correctly identify no connections when patterns are independent and uncorrelated.

#### 3.1.2 Scenario 2: Testing the methods’ ability to capture connectivity between two regions with a bivariate dependency between their patterns

Figure 5a shows how well the methods perform in the case of a bivariate, one-to-one dependency between two regions in the presence of other random patterns (Figure 2-2). The ground truth matrix highlights that the EV between *A*_1_ and *A*_2_ should be 1. The average EV for mvMDPC is 0.5, while the result of MDPC is 0.6. Figure 5b indicates the Euclidian distances of mvMDPC and MDPC on the left-hand side axis, as a function of SNR (on the x-axis), as well as the t-values for their comparison on the right-hand side y-axis. This curve suggests that for SNRs of zero and above, MDPC performs significantly better (indicated by black stars) than mvMDPC, and mvMDPC only performs significantly better when the SNR equals -10db (indicated by the green star, but the numerical difference is very small). Thus, for bivariate relationships the bivariate method performs better than the multivariate one. However, both are able to identify the bivariate dependency and easily distinguish it from the regions without a connection.

**Figure 5.**
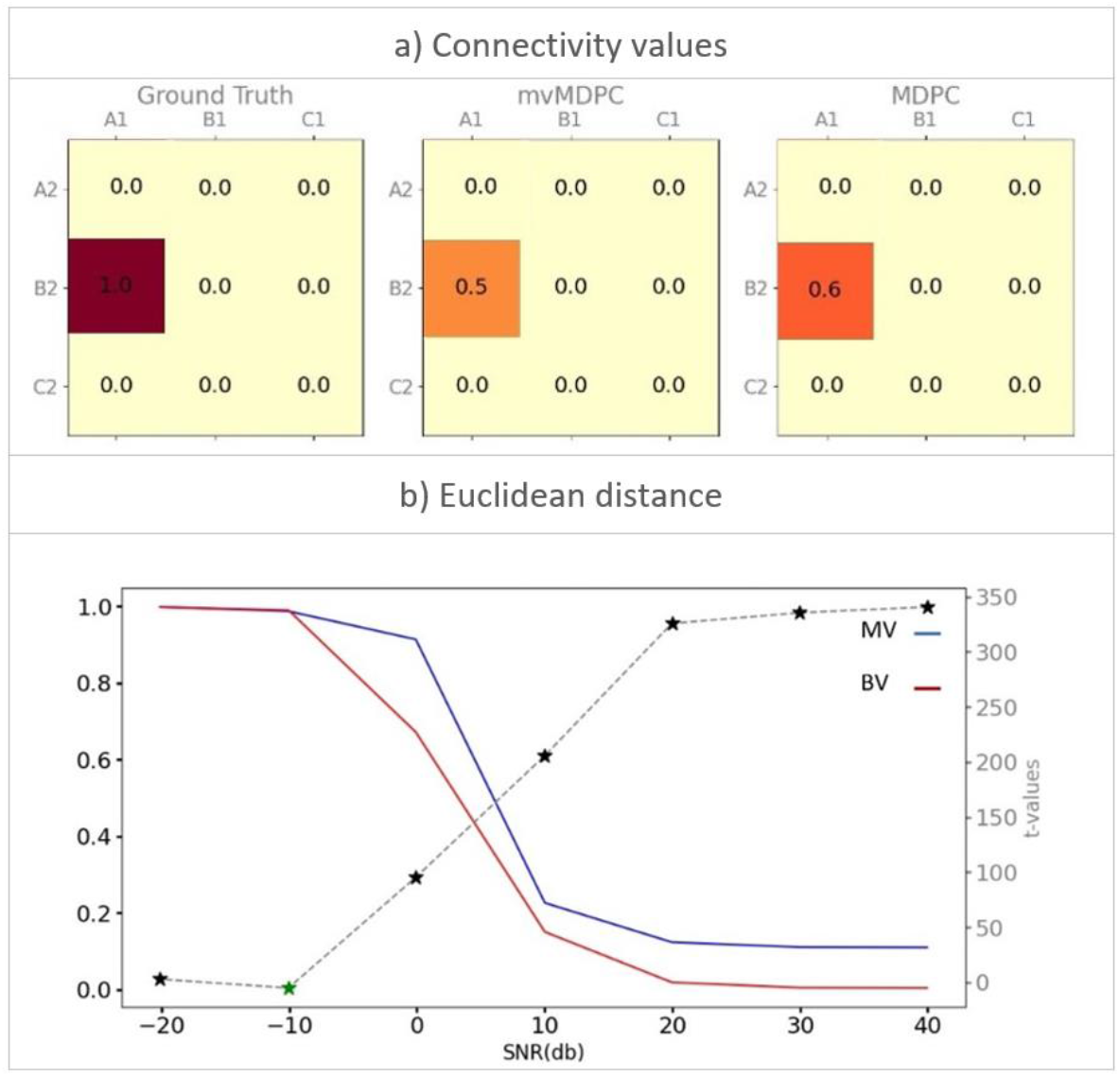
Representation of bivariate patterns. a) Connectivity values (explained variance) for the ground truth, mvMDPC, and MDPC methods (both averaged across SNRs). b) Euclidean distances for mvMDPC (blue curve) and MDPC (red curve) as a function of SNR as well as t-values for their direct comparison (dashed grey curve, scale on right-hand side y-axis). Black stars show where MDPC significantly outperforms mvMDPC, and green stars, when mvMDPC significantly outperforms MDPC. In almost all cases (except for -10db) the bivariate method performs significantly better.

#### 3.1.3 Scenario 3: Testing the sensitivity of each method to multivariate dependencies with different strengths

Figure 6a indicates the connectivity identified by each method when there are multiple influences with different strengths on one region. The ground truth highlights the greatest connectivity between *C*_1_-*A*_2_, then *B*_1_-*A*_2_, and finally *A*_1_-*A*_2_. While the averages generated by both methods reflect the decreasing strengths of connectivity correctly (i.e., decreasing EV from *C*_1_-*A*_2_, then *B*_1_-*A*_2_, then *A*_1_-*A*_2_), the multivariate method better captures the weaker relationships. As such the statistical comparison demonstrates that mvMDPC performs significantly better across almost all SNRs (green stars) when noise is present (see Figure 6b).

**Figure 6.**
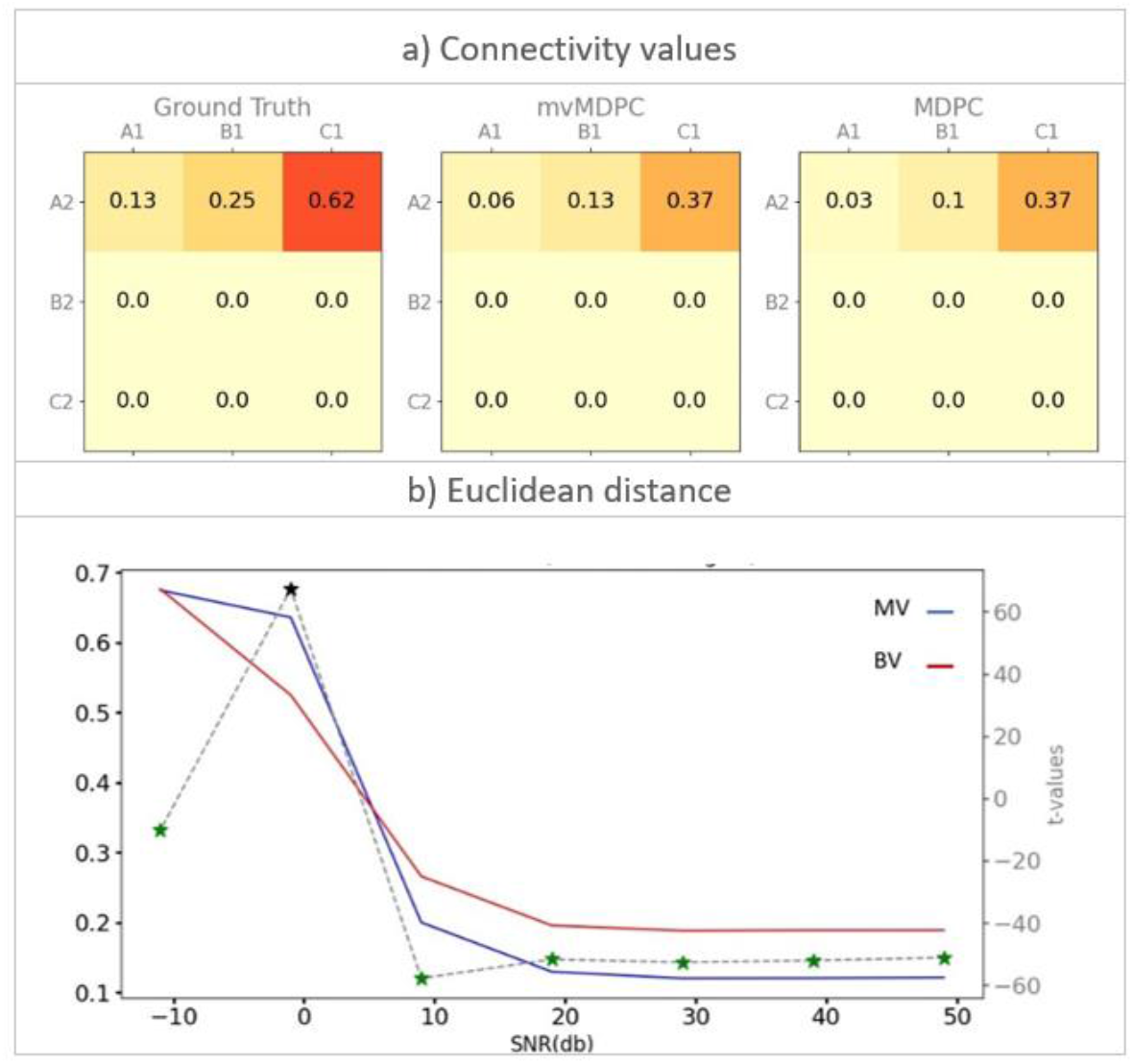
Simulation results for multivariate transformations with different strengths. a) Connectivity values (explained variance) for the ground truth, mvMDPC, and MDPC methods (both averaged across SNRs). b) Euclidean distances for mvMDPC (blue curve) and MDPC (red curve) as a function of SNR as well as t-values for their direct comparison (dashed grey curve, scale on right-hand side y-axis). Black stars show where MDPC significantly outperforms mvMDPC, and green stars when mvMDPC significantly outperforms MDPC. In all cases except for 0db, the multivariate method significantly outperforms the bivariate method.

#### 3.1.4 Scenario 4: Testing the ability of each method to capture multivariate all-to-all dependencies between patterns

Figure 7a shows the performance of each method when all-to-all, multivariate dependencies exist. Both methods are able to identify all connections in the simulated network. However, while the ground truth matrix suggests that the connectivity between all pairs should be 0.33, mvMDPC can only detect 0.17-0.18 of the explained variance across different SNRs and MDPC detects even less, 0.14-0.15. Figure 7b shows performance as a function of SNR. In almost all cases, mvMDPC significantly outperforms MDPC (indicated by green stars). Thus, for multivariate connectivity, the mvMDPC method produces more accurate outcomes.

**Figure 7.**
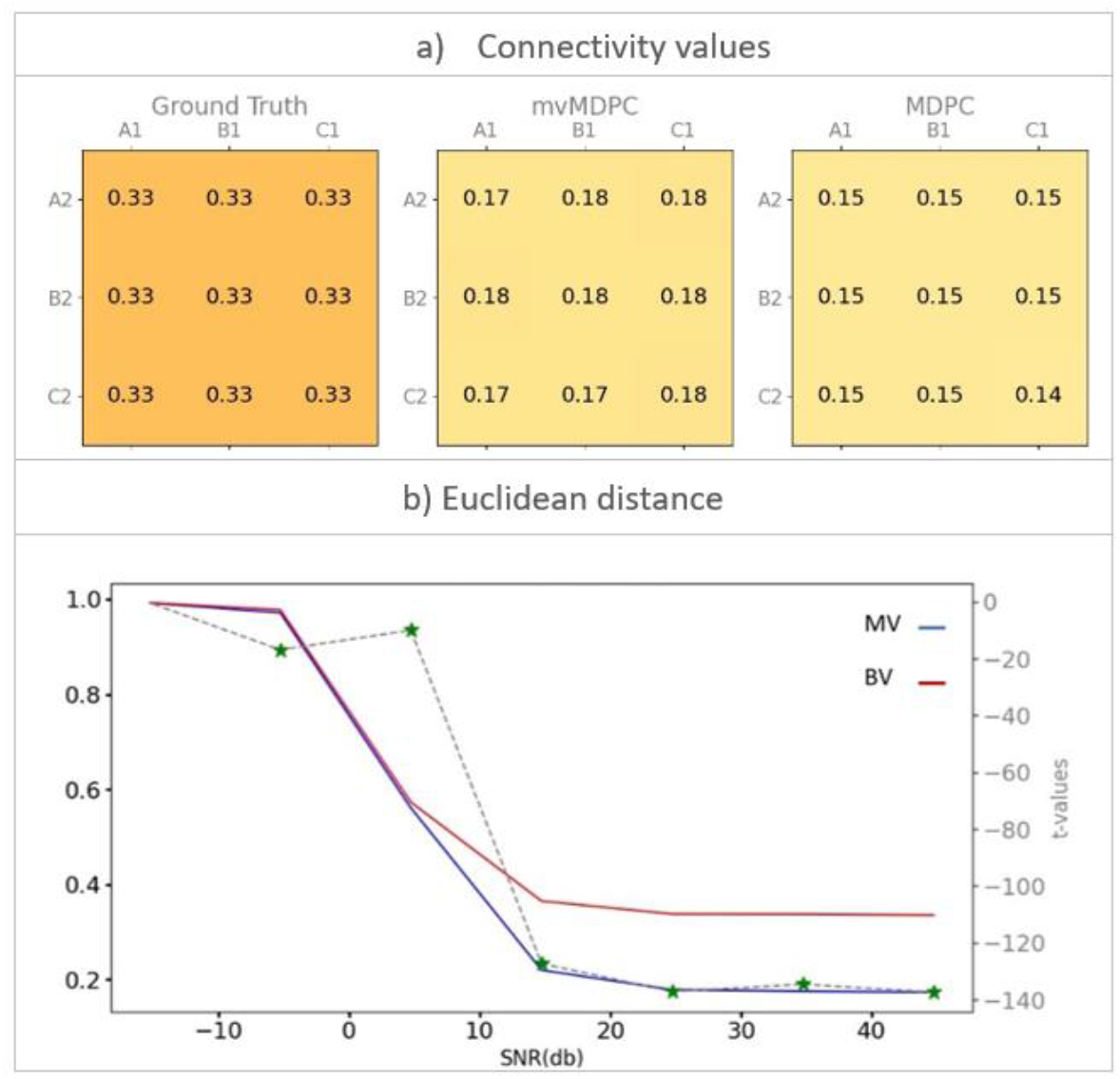
Simulation results for all-to-all, multivariate patterns. a) Connectivity values (explained variance) for the ground truth, mvMDPC, and MDPC methods (both averaged across SNRs). b) Euclidean distances for mvMDPC (blue curve) and MDPC (red curve) as a function of SNR as well as t-values for their direct comparison (dashed grey curve, scale on right-hand side y-axis). Black stars show where MDPC significantly outperforms mvMDPC, and green stars, when mvMDPC significantly outperforms MDPC. From around -5db onwards, the multivariate method significantly outperforms the bivariate method.

#### 3.1.5 Scenario 5: Testing the likelihood of each method to falsely identify indirect dependencies

We then tested how well the MDPC and mvMDPC methods perform in the case of an indirect dependency. Figure 8 shows that both methods detect the one true direct connection *B*_1_-*B*_2_ correctly, but both also capture the indirect connection *A*_1_-*B*_2_ to some extent. However, the mvMDPC approach estimates a much lower connectivity strength for this pair than for the true connection, significantly outperforming MDPC from -10db. The mvMPDC method produces more realistic EV values as it is less susceptible to this indirect connection and therefore better able to distinguish the direct and indirect connections.

**Figure 8.**
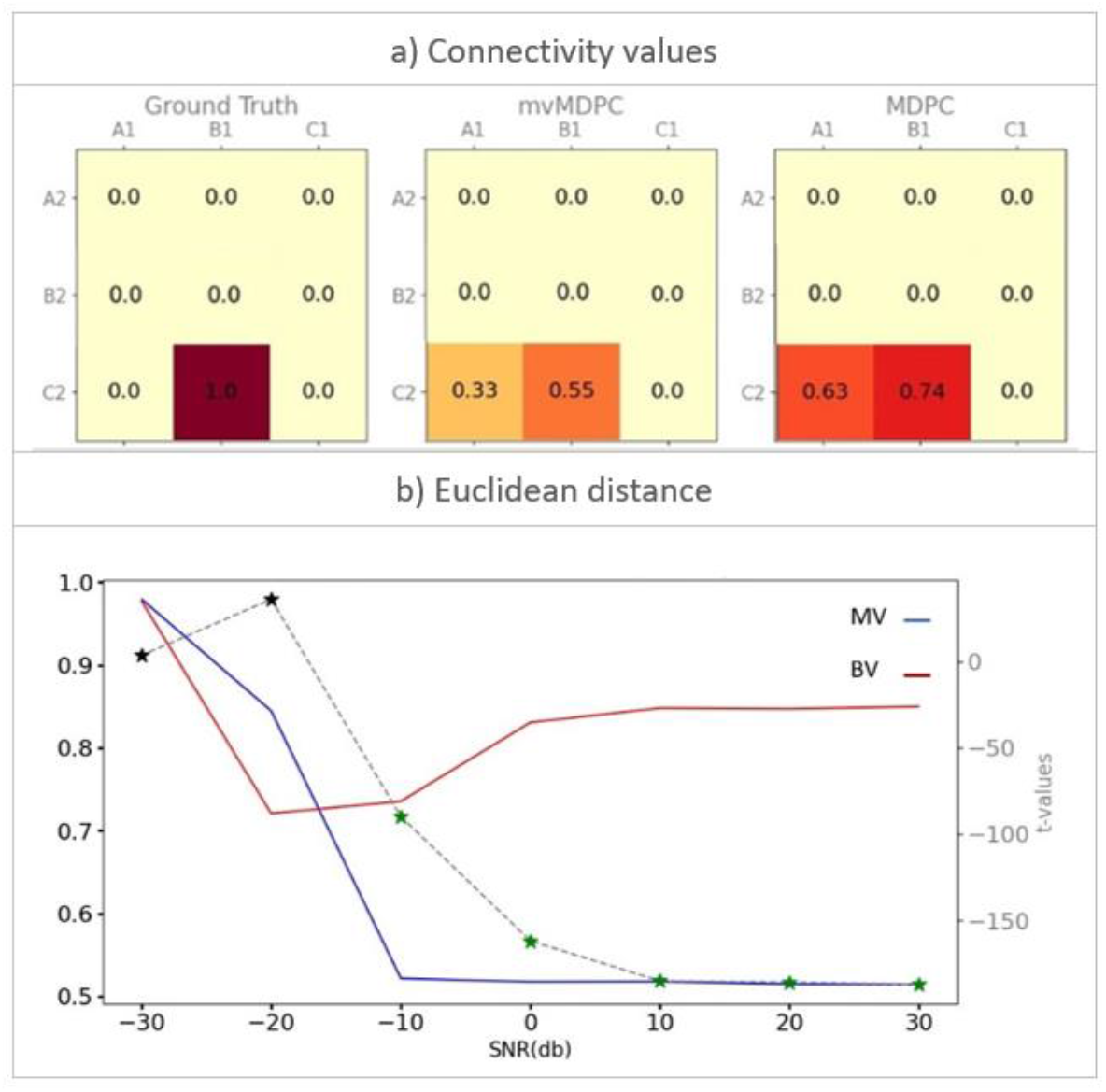
Simulation results for indirect dependencies. a) Connectivity values (explained variance) for the ground truth, mvMDPC, and MDPC methods (both averaged across SNRs). b) Euclidean distances for mvMDPC (blue curve) and MDPC (red curve) as a function of SNR as well as t-values for their direct comparison (dashed grey curve, scale on right-hand side y-axis). Black stars show where MDPC significantly outperforms mvMDPC, and green stars, when mvMDPC significantly outperforms MDPC. From around -10db onwards, the multivariate method significantly outperforms the bivariate method, while the bivariate method is significantly better below -10db.

#### 3.1.6 Scenario 6: Testing the likelihood of each method to spurious dependencies

We then tested how well the MDPC and mvMDPC methods perform in the case of a spurious dependency, where a connection may be falsely identified due to the shared influence of a third area. Figure 9a shows that both methods detect the direct connection *A*_1_-*C*_2_ correctly, yet both also capture the spurious connection *B*_1_-*C*_2_. Figure 9b indicates that the mvMDPC approach significantly outperforms the MDPC from 10db onwards as well as at -10db. The mvMDPC method is also doing somewhat better at distinguishing the real and indirect or spurious connections in terms of the ratio of their connection strengths. Overall, mvMDPC is better equipped than the bivariate method to distinguish true direct connections from spurious and indirect connections.

**Figure 9.**
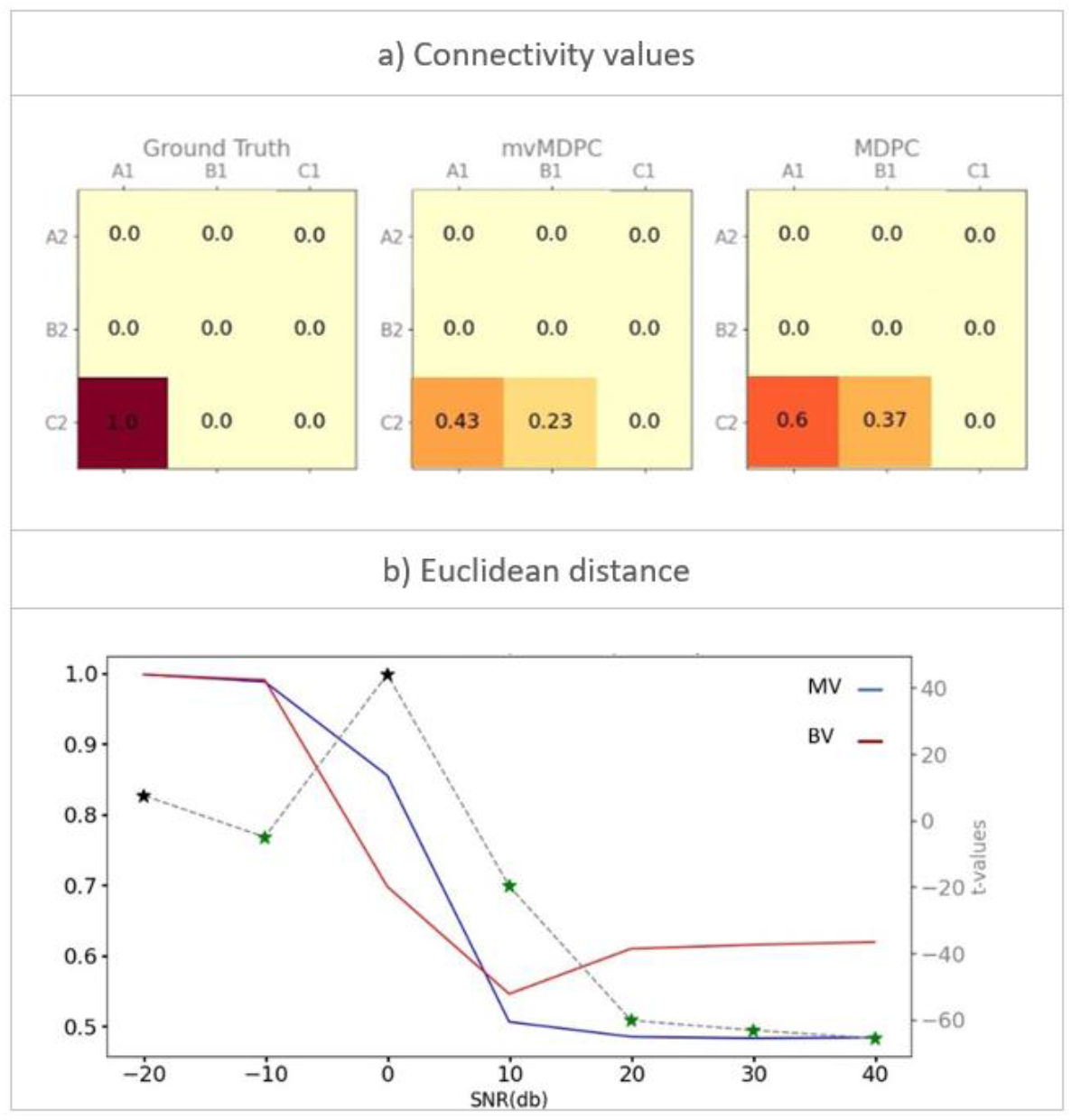
Representation of spurious dependencies. a) Connectivity values (explained variance) for the ground truth, mvMDPC, and MDPC methods. b) Euclidean distances for mvMDPC (blue curve) and MDPC (red curve) as a function of SNR as well as t-values for their direct comparison (dashed grey curve, scale on right-hand side y-axis). Black stars show where MDPC significantly outperforms mvMDPC, and green stars, when mvMDPC significantly outperforms MDPC. From around 10db onwards as well as at -10db, the multivariate method significantly outperforms the bivariate method.

#### 3.1.7 Random network simulations

Finally, we performed a more exhaustive comparison across multiple scenarios by running simulations across thousands of random networks and compared the Euclidean distance between the connections estimated by each method and the ground truth. The paired t-tests revealed significant differences in the performance of the two methods under different network conditions: without zero-lag dependencies: t-value = −35.23, p-value < 0.0001 and with zero-lag dependencies: t-value = −44.19, p-value < 0.0001. Regardless of whether zero-lagged dependencies are present, the mvMDPC method produced more accurate results, performing significantly better than MDPC. Overall, by considering the effect of all regions in one model, the multivariate method yields more reliable and realistic results.

Figure 10 shows the performance of the two methods as a function of SNR. Without zero-lag dependencies both bivariate and multivariate methods perform similarly for SNRs below zero, while both methods produce similar results for SNRs above zero, albeit with slightly lower Euclidean distances for the multivariate method. For networks with zero-lag dependencies, as shown in Figure 10b, the pattern is similar for negative SNRs. However, the bivariate method generates more variable Euclidean distances for positive SNRs, while those of the multivariate method are more focused around zero. The multivariate method therefore demonstrates greater consistency and reliability. Overall, the significant differences between the two methods are explained by the better performance of the multivariate method, especially when zero-lag dependencies are present. Note that zero-lag dependencies are expected to be present in real EEG/MEG data, both due to leakage and due to the effects of connections at prior times.

**Figure 10.**
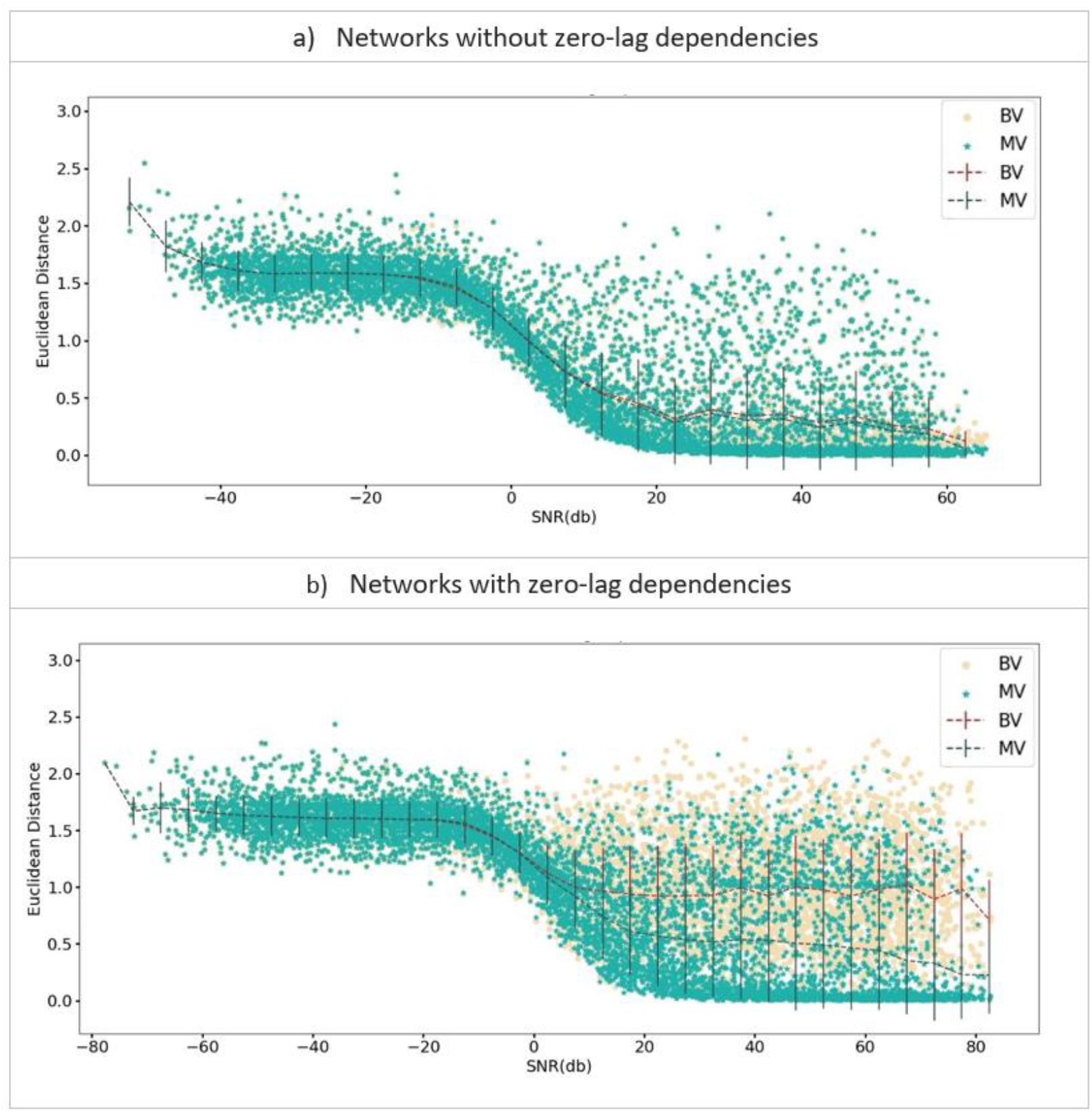
Scatter plots of Euclidean distances for simulated networks with random connections, comparing bivariate (yellow) and multivariate (green) methods. Dashed lines represent the average values in 10-unit intervals. a) Euclidean distance for networks without zero-lag dependencies plotted as a function of Signal-to-Noise Ratio (SNR); b) similar to a), but for networks with zero-lag dependencies. Both plots indicate a decline Euclidean distance measurements as SNR increases. While both methods struggle with high noise levels, the bivariate method shows greater variability in positive SNRs, especially when zero-lag dependencies are present. In contrast, the multivariate method is more consistent and reliably identifies connectivity patterns close to the ground truth.

### 3.2 Comparing mvTL-MDPC and TL-MDPC through application to a real EEG/MEG dataset

The simulations demonstrate that the multivariate method yields more accurate and reliable results across a range of network structures, detecting fewer spurious and indirect connections and thus better highlighting the true direct connections driving the network dynamics. Previously we applied the bivariate method to an EEG/MEG dataset comparing a semantic decision task with a lexical decision task (Farahibozorg, 2018; Rahimi et al., 2022). This revealed a highly connected semantic network, with most regions connected at a wide range of time lags. Our simulation results suggest this could be overinclusive, both across regions and times, as the bivariate method is insensitive to the difference between true direct connections and spurious or indirect connections. Even connections that are present at one time lag can show indirect or spurious connections at other lags as the shared past influence on the regions results in a zero-lag connection. If these results are overinclusive due to the bivariate method detecting spurious and indirect connections, the results may change with the current multivariate method, whereas if these are all true direct connections they should be detected by both methods. To distinguish these possibilities and gain a more accurate estimate of the network connectivity involved in controlled semantic cognition, we applied mvTL-MDPC to the same EEG/MEG dataset and compared the results to those obtained with the bivariate versions (Farahibozorg, 2018; Rahimi et al., 2022). Specifically, we asked if and how the connectivity of the semantic network depends on task demands, and whether mvTL-MDPC captures different information regarding the dynamic influences across this network than its bivariate version, TL-MDPC.

To guide interpretation of the full results, Figure 11 illustrates the outcome of the TL-MDPC (panel a) and mvTL-MDPC analyses (panel b) in the form of TTMs demonstrating the connectivity between lATL and IFG over time. These regions are critical for semantic representation and control, respectively. For both panels, the left TTM shows the average connectivity for the SD task across participants, the middle TTM shows the same for the LD task, and the right TTM reflects the statistical contrast (using cluster-based permutation tests) between the two tasks. An entry on the (*x,y*) coordinate of the TTMs describes the relationship between patterns in IFG at time point *x* and lATL at time point *y*. Note that neither method is strictly directional as a statistical relationship between patterns does not imply causality. However, where the activity patterns in one region predict those in another at a later time point, the order of the regions may be informative, suggesting one region is a leader and the other the follower. Not surprisingly, with both methods we found greater connectivity between these two areas for the task requiring deeper semantic processing, SD, compared to LD. This was true for all regions showing connectivity modulations and the remainder of the results focus on this between task comparison. One noticeable difference between the methods is that connectivity values are generally greater using the bivariate method; here the maximum is about 0.25, while that of the multivariate method is smaller, around 0.16. This likely reflects the fact that the multivariate method partials out variance from other regions. Importantly, as expected mvTL-MDPC shows significant task modulations in the connectivity of lATL and IFG at a more restricted set of time lags. This suggests that the areas found by both methods could reflect true direct influences between these areas. These connections (perhaps in combination with changes elsewhere in the network) may have led to spurious and indirect connections at the other time lags which were subsequently detected by TL-MDPC only. Note that the task modulations in connectivity identified with both methods are predominantly focused around the diagonal (reflecting a greater likelihood of instantaneous and relatively short lags than longer lags of influence). However, for TL-MDPC these effects were highly expansive (suggesting influences over hundreds of milliseconds) and were quasi-symmetrical, including lags on both sides of the diagonal to a similar extent. In contrast, mvTL-MDPC identified connectivity modulations much closer to the diagonal (with most influences occurring within a hundred milliseconds), as well as additional longer lags in the upper diagonal of the matrices, resulting in less symmetry. Thus, the mvTL-MDPC analysis suggests that statistical dependencies between lATL and IFG activity exist at short lags and are otherwise typically found between past patterns of IFG and future patterns of lATL. This provides additional information compared to the TL-MDPC analysis. Furthermore, it suggests that connectivity at additional time lags may be identified when using bivariate methods, yet this may be artefactual.

**Figure 11.**
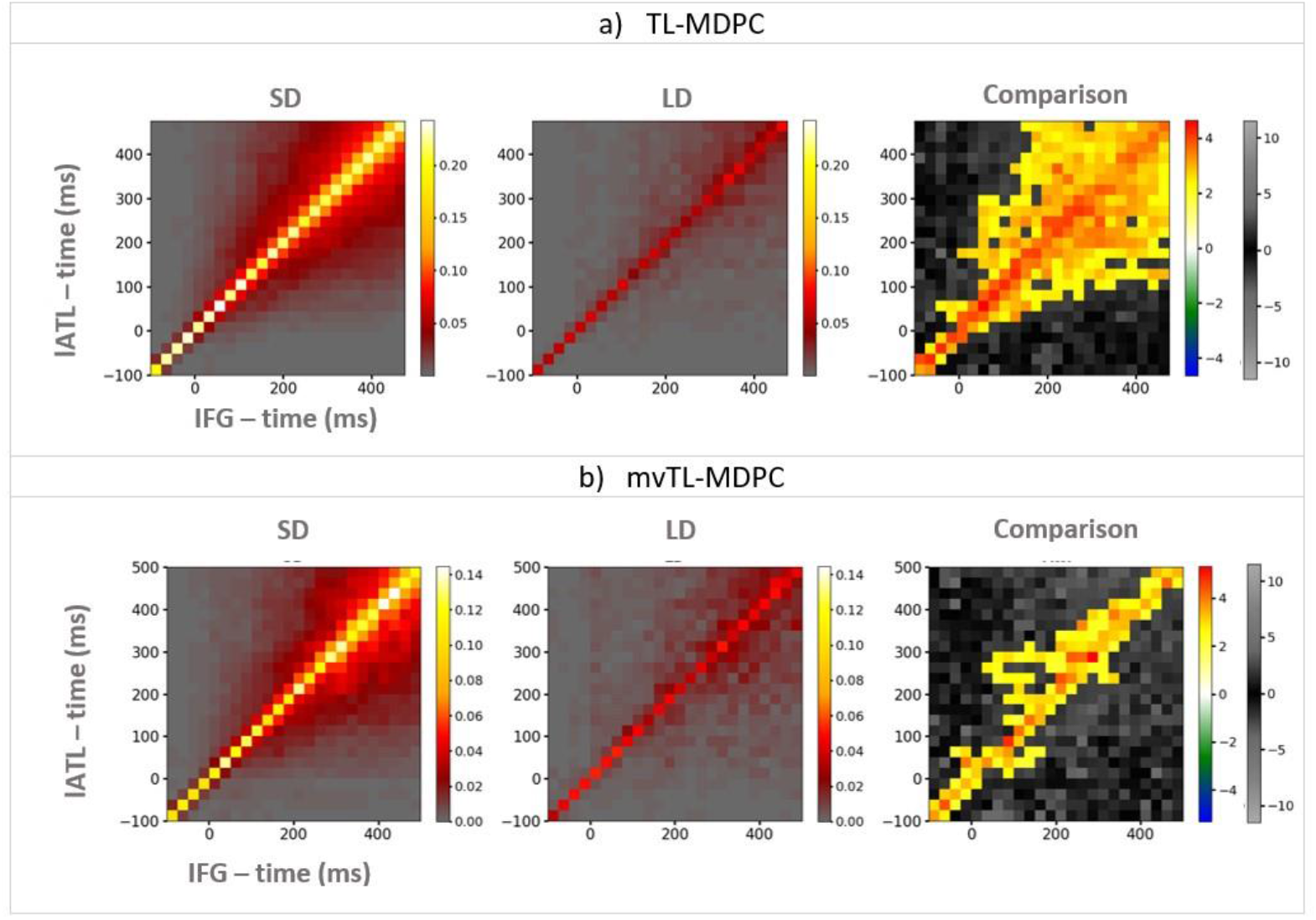
An illustration of TTMs showing the connectivity between lATL (y-axis) and IFG (x-axis), for the semantic decision (SD) task (left column), the lexical decision (LD) task (middle column), and their statistical contrast (right column). a) TTMs for TL-MDPC, b) TTMs for mvTL-MDPC. For both methods, the greatest connectivity is obtained around the diagonal. However, the statistical comparison between the tasks highlighted connectivity modulations at a large number of time lags on either side of the diagonal (symmetrically) using TL-MDPC. In contrast, mvTL-MDPC identified more circumscribed modulations particularly at shorter time lags. Additionally, significant increases in connectivity were identified more frequently between past patterns of IFG and future patterns of lATL than the other way around. The simulations above suggest that mvTL-MDPC is more reliable and the additional connections shown only with the bivariate method may be due to indirect and spurious dependencies. Thus, lATL and IFG may not directly connect at these time lags. Colour bars show connectivity scores (explained variance) for the first two columns. For the third column, the first colour bar (hot and cold colours) highlights significant effects based on the cluster-based permutation test, while the grey-scale colour bar indicates non-significant t-values (this colour bar is the same for all columns).

#### 3.2.2 Capturing the connectivity within the semantic network across time with mvTL-MDPC

We computed the connectivity between all 15 pairs of ROIs using TL-MDPC and mvTL-MDPC, and present the resulting inter-regional connectivity matrix (ICM) (Rahimi et al., 2023a) in Figure 12. Here we focus on the contrast between the two tasks, but TTMs of each task and each pair of regions are presented in Figure S1 in the Supplementary Materials. Figure 12 illustrates the statistical comparison between the semantic and lexical decision tasks as TTMs for each pair of ROIs, with the upper diagonal (yellow area) showing the outcome of mvTL-MDPC, and the lower diagonal (blue area) representing the TL-MDPC results. As the results differed by direction for the multivariate method, the values were only estimated in a single (past to future) direction, providing a second difference from the bivariate method in addition to the use of all the ROIs to estimate a single model. To check that this was not the cause of the differences between the two methods we also ran the bivariate method in a single (past to future) direction, shown in Figure S2 of the Supplementary. The results were highly similar to the bidirectional version and the differences to the multivariate method remained. Therefore, this factor cannot explain the differences between the multivariate and bivariate methods.

**Figure 12.**
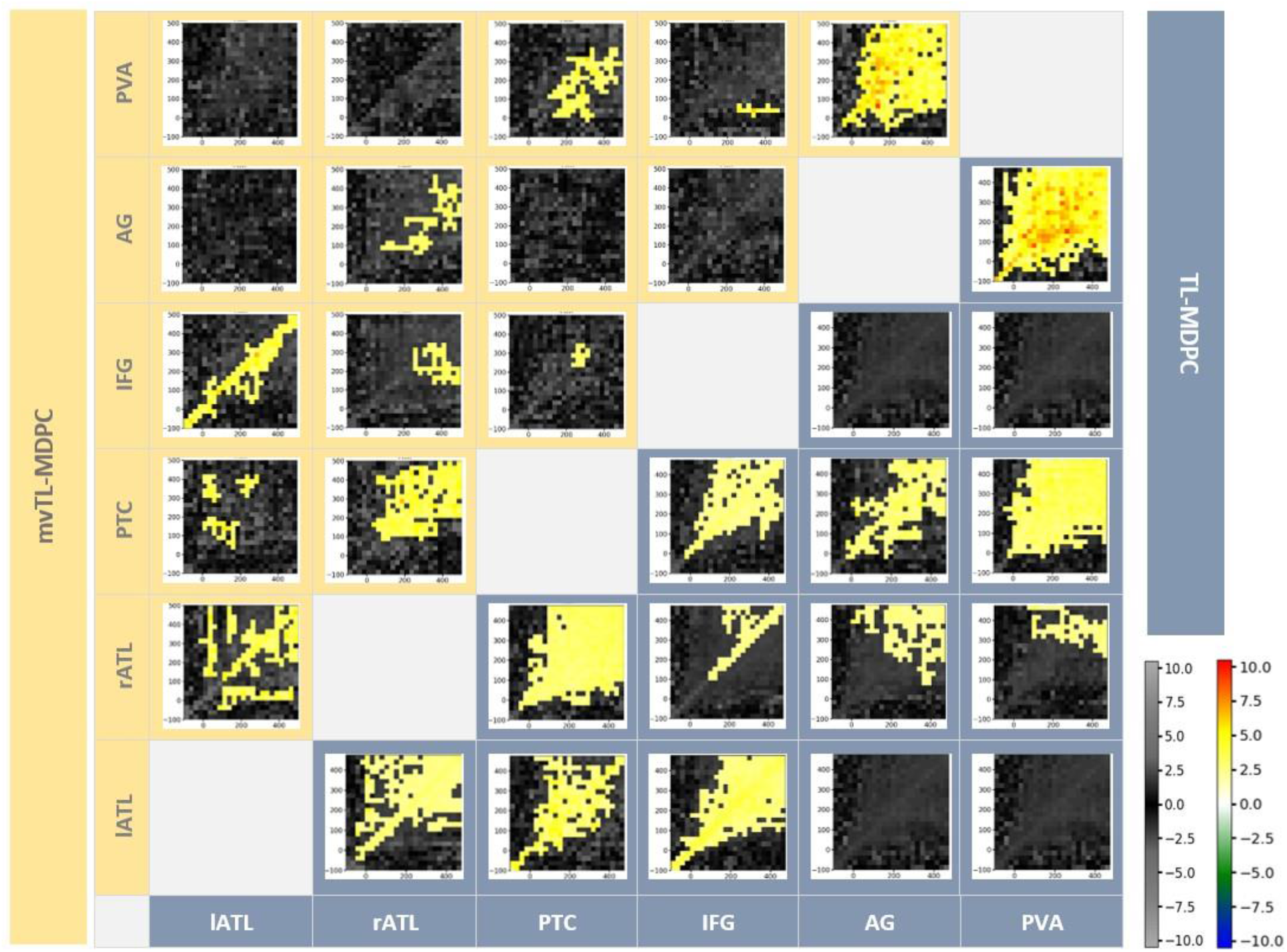
Inter-regional Connectivity Matrix (ICM) for semantic network modulations in the brain. The upper diagonal (yellow shaded area) shows the mvTL-MDPC results and the lower diagonal (blue shaded area) shows the TL-MDPC results. For each method there is a TTM showing connectivity (explained variance) across time lags for each pair of regions. All significant modulations obtained by both methods revealed more connectivity for SD than LD. Cluster size was thresholded at 2% of TTMs size (24*24). Both methods detected rich modulations between semantic control and representation regions. As expected the connections identified with mvTL-MDPC were more circumscribed than for TL-MDPC, with fewer pairs showing connectivity across fewer time lags. The colour bar on the right (hot and cold colours) highlights significant effects based on the cluster-based permutation test, while the grey-scale colourbar indicates non-significant t-values for all TTMs.

TL-MDPC identified extensive connectivity modulations across 11 pairs of ROIs, each showing greater connectivity for the more semantically demanding task. Most of these pairs, including lATL with rATL, PTC, and IFG, as well as rATL with PTC, PTC with IFG and AG with PVA, demonstrate connectivity changes over many time lags, fanning out from the diagonal and including many connections present over very long time lags (for instance, there are dependencies between PTC and PVA activity patterns over 400ms apart). Connectivity between rATL and IFG occurred at later time points and was more circumscribed, mostly confined to the diagonal. In addition, there were late, symmetrical effects of connectivity between rATL and both AG and PVA, indicating connectivity across long time lags but only at later time points. mvTL-MDPC also identified connections between nine of these ROI pairs, supporting their role in controlled semantic processing. However, it did not identify significantly greater connectivity modulations between rATL and PVA or PTC and AG. These may therefore reflect spurious or indirect connections that are falsely detected by bivariate methods. Moreover, whilst connectivity was still identified between most of the pairs of regions previously highlighted, the time lags showing significant connections between each of these pair of ROIs were far more circumscribed (albeit with a fairly extensive connection remaining between AG and PVA). For the connections between lATL and rATL, rATL and PTC and AG and PVA this reduction did little to change the pattern of significant connections across time lags. These connections showed a symmetrical pattern across a sustained period, possibly reflecting the existence of continued, recurrent activation flow between these regions. lATL and rATL and AG and PVA showed significant connectivity throughout, with connectivity over long time lags. rATL and PTC showed a similar pattern of effects over long time lags, with significant connectivity modulations beginning around 100ms after word onset.

For all other pairs of connected regions, the multivariate method resulted in a change in the distribution of the identified connections to an asymmetrical pattern, reflecting more evidence for past patterns of one ROI predicting future patterns of the other ROI than the vice versa. For example, past patterns of PVA predict future patterns of PTC and IFG, suggesting information flow from early visual to higher level processing regions. As described above, there is also evidence for connectivity between lATL and IFG throughout, with significant predictions over short lags and a greater likelihood of past IFG predicting future lATL. At later times, IFG also predicts future rATL patterns, and past patterns of lATL predict future patterns of PTC. PTC and IFG, the two putative control regions, predict each other over short lags at later time periods. Past patterns in AG predict future patterns of rATL.

In addition, mvTL-MDPC also highlighted connectivity between IFG and PVA which was not identified by the bivariate method. Despite the small size of this cluster, its existence is plausible, considering the temporal dynamics between visual and higher-order semantic areas, as well as the direction of connectivity. Here PVA patterns present shortly after onset predicted late IFG patterns around 400ms. In general, the multivariate method produced significant connectivity between most, yet not all, of the same regions and at far fewer latencies and shorter time lags, with less symmetrical TTMs highlighting the temporal order of the related activity patterns in the connected areas. This is highly compatible with the simulation results, suggesting that the multivariate method is less likely to capture spurious and indirect connectivity, particularly over time, and therefore produces a more reliable and informative picture of the semantic network.

Figure 13 shows the dynamic brain semantic network obtained from the mvTL-MDPC at 100, 200, 300, 400, and 500ms. This figure reflects the sum of significant t-values (*min*=2, *max*=42) at the specified time points and their corresponding lags. For instance, the arrow from lATL to rATL at 200ms represents the sum of all significant t-values from the past of lATL (before 200ms up to 200ms) to 200ms at rATL. For visualisation purposes, the sum values are categorised into eight groups, represented by the width of the arrows, with thicker arrows indicating higher values. The figure illustrates how the connectivity strength varies over time and lags for all pairs. With the multivariate method, the ATLs remain highly connected to other areas of the semantic network, while connectivity of the AG is sparser, with only connections with PVA and rATL.

**Figure 13.**
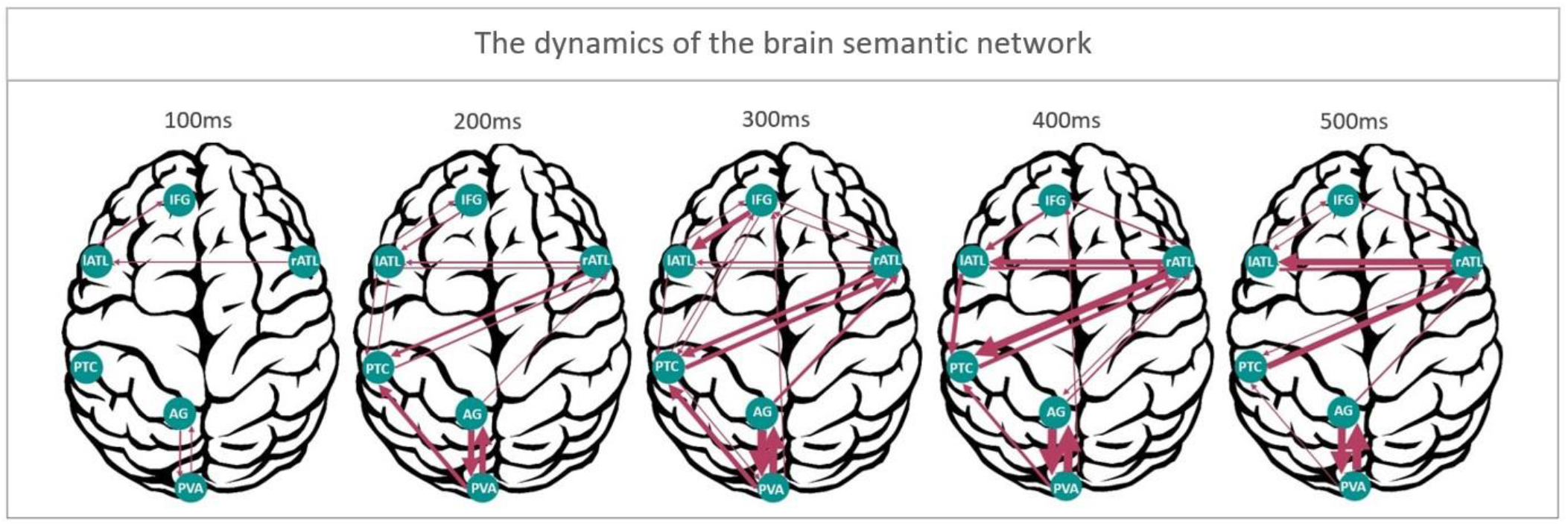
Representation of the dynamics of the cortical semantic network. Connectivity strength, highlighted using the mvTL-MDPC method, is shown at 5 different time points (100, 200, 300, 400, and 500ms). This illustrates how the connectivity strength varies over time and lags for all pairs by reflecting the sum of significant t-values at the specified time points and their corresponding lags. For instance, the arrow from lATL to rATL at 200ms represents the sum of all significant t-values from the past of lATL (before 200ms) up to 200ms at rATL. For visualisation purposes, the summed t-values are categorised into eight equally-sized groups, represented by the width of the arrows, with thicker arrows indicating higher values.

## 4 Discussion

We introduced a novel functional connectivity method for event-related EEG/MEG data, multivariate time-lagged multidimensional pattern connectivity (mvTL-MDPC). This method is designed to detect linear relationships between activation patterns within pairs of regions as they unfold over the time course of a trial, while accounting for the contributions of multiple other regions. To validate this method and compare it with the bivariate version, TL-MDPC, we conducted exhaustive simulations on both structured and random networks. These simulations indicated that the multivariate method is more reliable, yielding results that are closer to the ground truth in the majority of cases. Accounting for the impact of other regions, improved the method’s ability to distinguish direct connections from indirect or spurious effects caused by the relationship of the ROIs with additional areas, or with each other at different time points. Thus, we expected the connections identified in real EEG/MEG data to be more selective when using this multivariate method. Indeed, in our real EEG/MEG application, mvTL-MDPC resulted in fewer connections compared to TL-MDPC, with both fewer regions being demonstrated to connect, and regions connecting at fewer time lags. Despite this, mvTL-MDPC still effectively captured the connectivity between core semantic regions, suggesting it provides more accurate discrimination of direct connections from indirect and spurious connections. As a result of this improved discrimination ability, the results were less symmetrical, with the activity pattern in one region more clearly following the activity pattern in the other. Thus, while still being a functional rather than effective connectivity method, it provides additional information to judge the temporal order of the connections compared to bivariate analyses, improving our ability to determine how information is transmitted across task networks.

Our simulations showed that the multivariate method outperforms the bivariate method in a range of complex scenarios. In particular, the multivariate method was less prone to identifying spurious and indirect connections, making it more robust. We observed this pattern of results for multivariate transformations with different strengths, all-to-all multivariate mappings, indirect dependencies, and spurious dependencies. As bivariate methods are likely to identify relationships that do not reflect a direct connection (Haufe et al., 2010), they may provide highly over-inclusive results, such as the detection of all possible connections within a task network. In contrast, the multivariate method is better able to protect against this, leading to more selective and therefore better interpretable findings. Our extensive comparisons of the two methods, involving simulations of thousands of random networks, demonstrated that mvTL-MDPC consistently outperforms TL-MDPC in accurately estimating network connections, irrespective of the presence of zero-lag dependencies. Furthermore, the multivariate method was more reliable in its ability to consistently detect the true connectivity pattern across iterations. Overall, the findings from our simulations suggest that the multivariate method offers greater reliability, discriminability, and interpretability than the bivariate approach.

The results of our real EEG/MEG data analysis support this conclusion. We confirmed that the multivariate method identified fewer connections than the bivariate method in EEG/MEG data. Additionally, most relationships identified with mvTL-MDPC occurred over a shorter timeframe, typically reflecting relatively brief relationships unfolding within a hundred milliseconds. This finding contrasts with the more expansive effects observed in TL-MDPC, which often spanned hundreds of milliseconds. These results imply that some of our previous findings (Rahimi et al., 2023a, 2023b) may reflect indirect or spurious connectivity. Indeed, the systematic preservation of connections over shorter time lags and reduction in connections over longer delays is highly consistent with the conclusion that the multivariate method is selectively identifying direct (and therefore fast-acting) relationships while removing indirect or spurious connections. Of course, it cannot be concluded that no indirect or spurious connections will be falsely identified using the multivariate method. However, there are likely to be fewer, reflecting an improvement on bivariate methods. Moreover, while the multivariate method could be failing to identify some true direct connections, this is not the key pattern of change demonstrated by the simulations, where the key differences related to the spurious and indirect relationships. Overall, the connections detected by the multivariate method are more likely to be reliable and realistic.

Importantly, our novel multivariate method corroborates the major findings of our previous study (Rahimi et al., 2023a) and is consistent with the Controlled Semantic Cognition framework (Jefferies, 2013; Lambon Ralph et al., 2016). In all of our analyses we observed rich connectivity for the ATLs, in contrast to relatively sparse connectivity for AG, a pattern that is more pronounced with the more selective multivariate method. These findings are consistent with the ATLs’ critical role as a hub for multimodal semantic representation (Lambon Ralph et al., 2016; Patterson et al., 2007; Rogers et al., 2006). Indeed, both left and right ATL demonstrated rich connectivity with regions throughout the semantic network, including each other. There is clear evidence for influences in both directions between left and right ATL despite the increased selectivity of the multivariate method, although at later latencies the influence of rATL on lATL appears stronger. This highlights the importance of both ATLs working in tandem.

It may initially appear surprising that the identified connections with other semantic regions are typically stronger for rATL than its left counterpart, despite the use of written verbal stimuli (Rice et al., 2015). Note, however, that the findings reflect the differential connectivity of two tasks with varying levels of semantic demands.. Specifically, the left ATL is highly engaged and connected with other semantic areas for both tasks, while the right ATL demonstrates a greater increase in engagement as the semantic demands increase, precisely because it is not automatically recruited when demands are low.

In contrast to the ATLs, the AG appears isolated from this semantic network, with only minimal connectivity to one semantic region, rATL. The connections observed between AG and semantic regions with the bivariate method may reflect indirect or spurious connectivity, perhaps due to their shared connectivity with visual input regions. Indeed, despite the apparent separation from the semantic network, the AG exhibited strong connectivity with the PVA. While its role is still debated, this finding is compatible with a distinct role for the AG, involving processing of incoming stimuli in a way which does not require integration with the semantic network. This may reflect more domain-general processing that is not specific to semantic stimuli, such as the extraction of statistical regularities across time (Humphreys et al., 2021) or the construction of a scene (Irish et al., 2015; Ramanan et al., 2018b, 2018a; Wilson et al., 2020). Overall, our findings provide little evidence for a key role of AG in the semantic network, but it may be involved in parallel context integration, episodic memory or attentional processes (Cabeza, 2008; Cabeza et al., 2012; Chambers et al., 2004; Farahibozorg et al., 2022; Humphreys et al., 2021; Humphreys and Lambon Ralph, 2015; Noonan et al., 2012; Shimamura, 2011; Vilberg and Rugg, 2008; Wagner et al., 2005).

We demonstrate a high degree of connectivity between the bilateral ATLs, critical for semantic representation, and the PTC and IFG regions responsible for semantic control, in a demanding semantic task, as per the controlled semantic cognition (CSC) framework (Lambon Ralph et al., 2016). The multivariate method highlights the connectivity of both IFG and PTC with the ATLs, with significant influences in both directions for each pair. This includes strong evidence for the influence of lATL on PTC and IFG on rATL at later time points. Despite the remarkable similarity of the roles attributed to PTC and IFG in semantic control processing (Jackson, 2021; Jefferies, 2013; Jefferies and Lambon Ralph, 2006; Lambon Ralph et al., 2016), the multivariate method finds relatively little evidence of direct influences between these regions, with only a small cluster of significant influences over short lags around 300ms post stimulus onset. However, it should be noted that there are multiple functional regions within PTC, including both control regions and areas contributing to the gradual transition from occipital visual to anterior temporal semantic representations (e.g. Hodgson et al., 2023). Consequently, it is possible we may be missing the more subtle control-related effects due to the spatial resolution of MEG and the absence of a task comparison specifically designed to isolate semantic control processes. A more specific manipulation of semantic control should be a key consideration for future studies exploring the connectivity of the semantic network. Note that the multivariate method detected interactions between many crucial semantic control and representation regions, demonstrating that it is still sensitive to the presence of multiple dependencies among regions despite partialling out their common variance.

Interestingly, the TTMs obtained from the multivariate method are notably less symmetrical than those created with the bivariate method. Consequently, mvTL-MDPC provides more evidence of one ROI’s past patterns predicting future patterns of another ROI than the reverse, instead of merely demonstrating that past patterns of either ROI predict future patterns of the other. As such, mvTL-MDPC offers greater insight into the temporal order of the connections, which we did not observe for TL-MDPC. For instance, the TTM for the PTC and PVA suggests that past patterns within the visual area predict future patterns in PTC, with less evidence for predictions the other way around, in line with strong effects of early feedforward processing in the visual system (Marinkovic et al., 2003; Serre et al., 2007). A possible exception to this increasing asymmetry is the connection between AG and PVA, which remains strongly bidirectional and present over a wide range of latencies. These areas may be strongly interdependent as their event-related responses are highly similar (Rahimi et al., 2022). Additionally, left and right ATL have strong interconnectivity from around 100ms, consistent with the view that they are two parts of one multimodal semantic representation hub, with strong interhemispheric interdependence (Jung and Lambon Ralph, 2016; Rice et al., 2015). Many other connections have dominant directions (lATL to PTC, IFG to rATL, AG to rATL and to a lesser extent, IFG to lATL). However, all of these have evidence for some influence in the opposing direction, although not necessarily at the same point in time. This bidirectional pattern provides evidence for recurrent activation flow between semantic control and representation regions. Recurrence has previously been suggested as an essential part of semantic object processing at early processing stages by combining EEG/MEG and neural network modelling (Kietzmann et al., 2019).

Consistent with prior studies (e.g. Marinkovic et al., 2003) and the bivariate method (Rahimi et al., 2023a), the strongest evidence for interactions between representation and control regions was found at later time points, critical for semantic cognition, around 300 to 500ms after stimulus onset. However, we also observed changes in their interaction far earlier, including the connectivity of rATL-PTC from around 100ms and lATL-IFG from around stimuli presentation. This is consistent with the evidence of task effects on early word recognition processes in previous EEG/MEG studies (Chen et al., 2015, 2013) and intracortical EEG evidence for the impact of frontal control regions on early feedforward semantic processing (Tiesinga et al., 2023). Together the findings suggest that the retrieval of task-relevant aspects of a concept is an interactive process, requiring the interplay of representation and control regions throughout the time course of the semantic decision, aligning with current models of controlled semantic cognition (Giallanza et al., 2023; Hoffman et al., 2018; Jackson et al., 2021; Rogers et al., 2021).

The performance of any EEG/MEG source connectivity method depends on the spatial resolution of the source estimation procedure. Source leakage is an inevitable consequence of the non-uniqueness of the EEG/MEG inverse problem, which can affect all unidimensional and multidimensional analyses, and in particular connectivity studies (Colclough et al., 2015; Farahibozorg et al., 2022; Hauk et al., 2022; Palva et al., 2018). To alleviate this issue and optimise spatial resolution, we used combined EEG and MEG in the current study. Similar to our previous studies on the same dataset (Rahimi et al., 2023b, 2023a, 2022), here we present a quantitative description of our EEG/MEG leakage between regions with non-homogeneous activations. Our leakage assessment suggests that the areas with the strongest leakage are not the same as those identified as having largest connections strengths by the multivariate method. Therefore, our results cannot be solely attributed to leakage. A more comprehensive evaluation of leakage in multivariate, multidimensional scenarios should be carried out in future research.

In conclusion, our new multivariate and multidimensional approach for estimating the dependencies between activation patterns across space and time enabled us to capture more reliable and realistic results of dynamic brain connectivity in event-related EEG/MEG data. While mvTL-MDPC is an undirected functional connectivity method, meaning that we cannot directly infer the causality or direction of information flow, we can rely on the timing information to make inferences about directionality. Compared to bivariate methods, it can provide more insight into network interactions as it is more likely to identify influences at specific time lags and in a single direction. This method could be extended further to identify effective connectivity using the rationale of Granger Causality and autoregressive models (Granger, 1969; Hu et al., 2012; Seth, 2010). Note that it is also possible to compute nonlinear transformations between patterns. In a previous study, we only found small differences between linear and non-linear TL-MDPC in our analysis of real EEG/MEG data, while the non-linear method came at a much higher computational cost (Rahimi et al., 2023b). Future studies should investigate whether this is also the case for a nonlinear multivariate and multidimensional version of this method.

Eventually, these methods could also be implemented in the frequency or time-frequency domain to unravel how those representations are transformed across brain regions, latencies, and frequencies. Adopting a multivariate, multidimensional approach has allowed a more detailed investigation of the semantic network across space and time compared to previous unidimensional and bivariate analyses using fMRI (Chiou et al., 2018; Jackson et al., 2016; Jung and Lambon Ralph, 2016) and EEG/MEG (Farahibozorg et al., 2022; Rahimi et al., 2023a, 2023b, 2022; Sormaz et al., 2017). We hope that this study is a useful step towards future methods development and research that can indeed “transform” spatiotemporal connectivity analyses.

## 5 Acknowledgements

This work was supported by intramural funding from the Medical Research Council UK (MC_UU_00005/18), a British Academy Postdoctoral Fellowship awarded to R.L.J. (no. pf170068), and Cambridge University international scholarship awarded to S.R. For the purpose of open access, the author has applied a CC BY public copyright license to any Author Accepted Manuscript version arising from this submission. We also thank Dr Rezvan Farahibozorg for data collection and information on the dataset.

## Conflicts of interest

The authors declare no conflicts of interest

## Supplementary Materials

**Figure S1.**
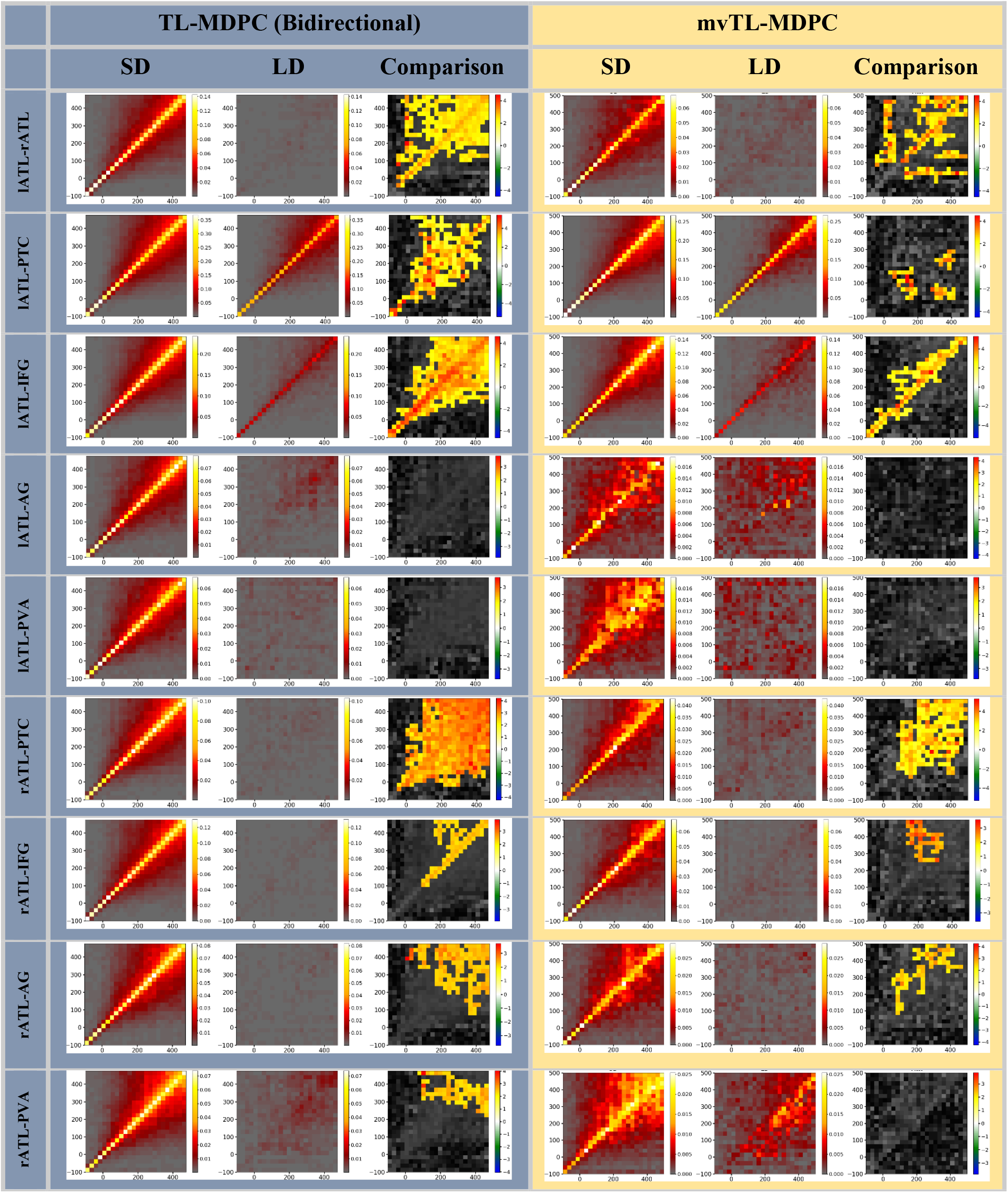
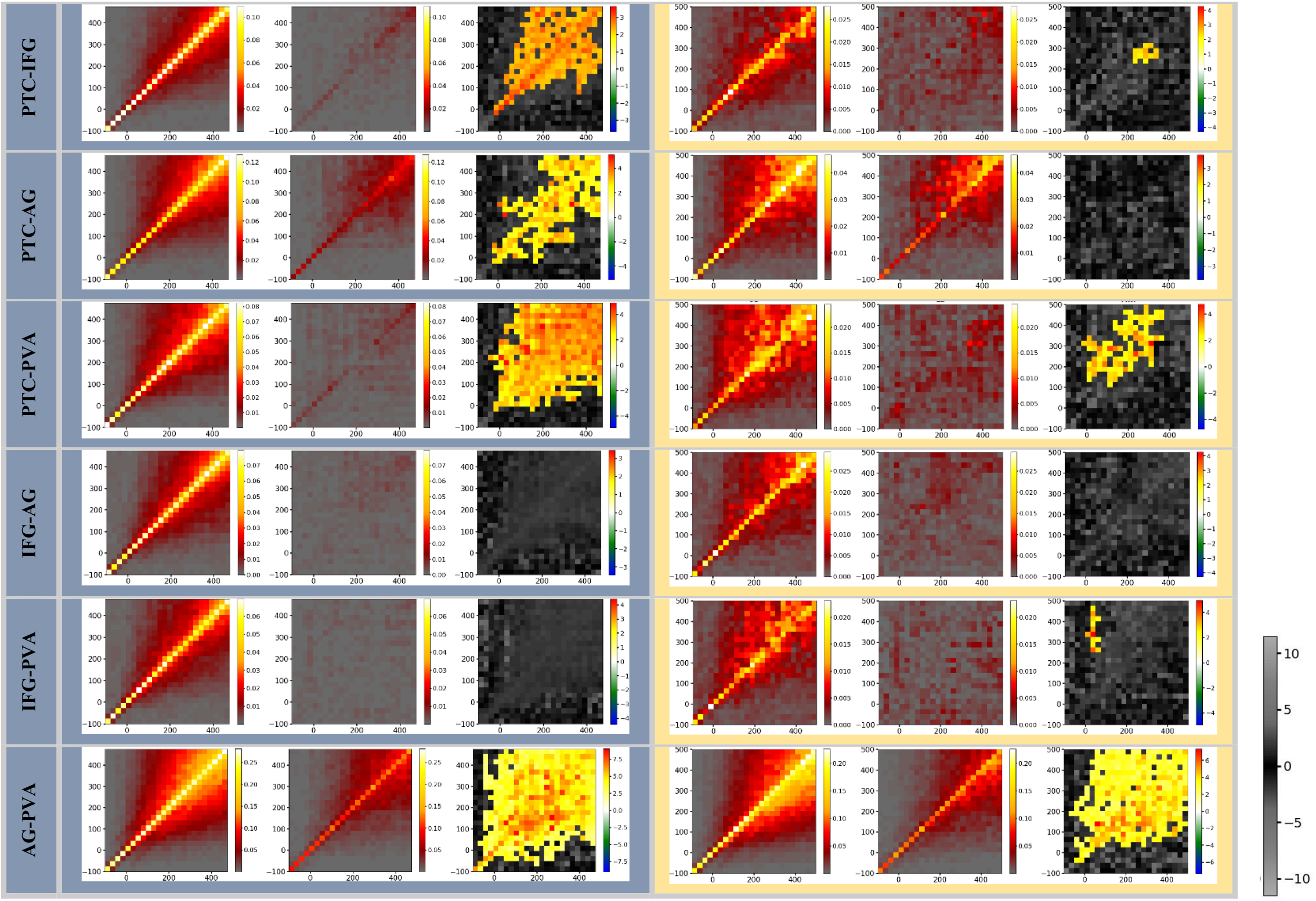
Representation of all TTMs for the semantic decision (SD) and lexical decision (LD) tasks, and their comparison, using TL-MDPC (left column, as computed in 2023a) and mvTL-MDPC (right column). TTMs for SD and LD are averaged across participants, and comparison was performed using cluster-based permutation test with alpha-level=0.05. All significant contrasts show greater connectivity for SD than LD. To be displayed here, the clusters must be larger than 2% of the TTMs size (24*24). The grey-scale colourbar indicates non-significant t-values (this colour bar is the same across all TTMs).

**Figure S2.**
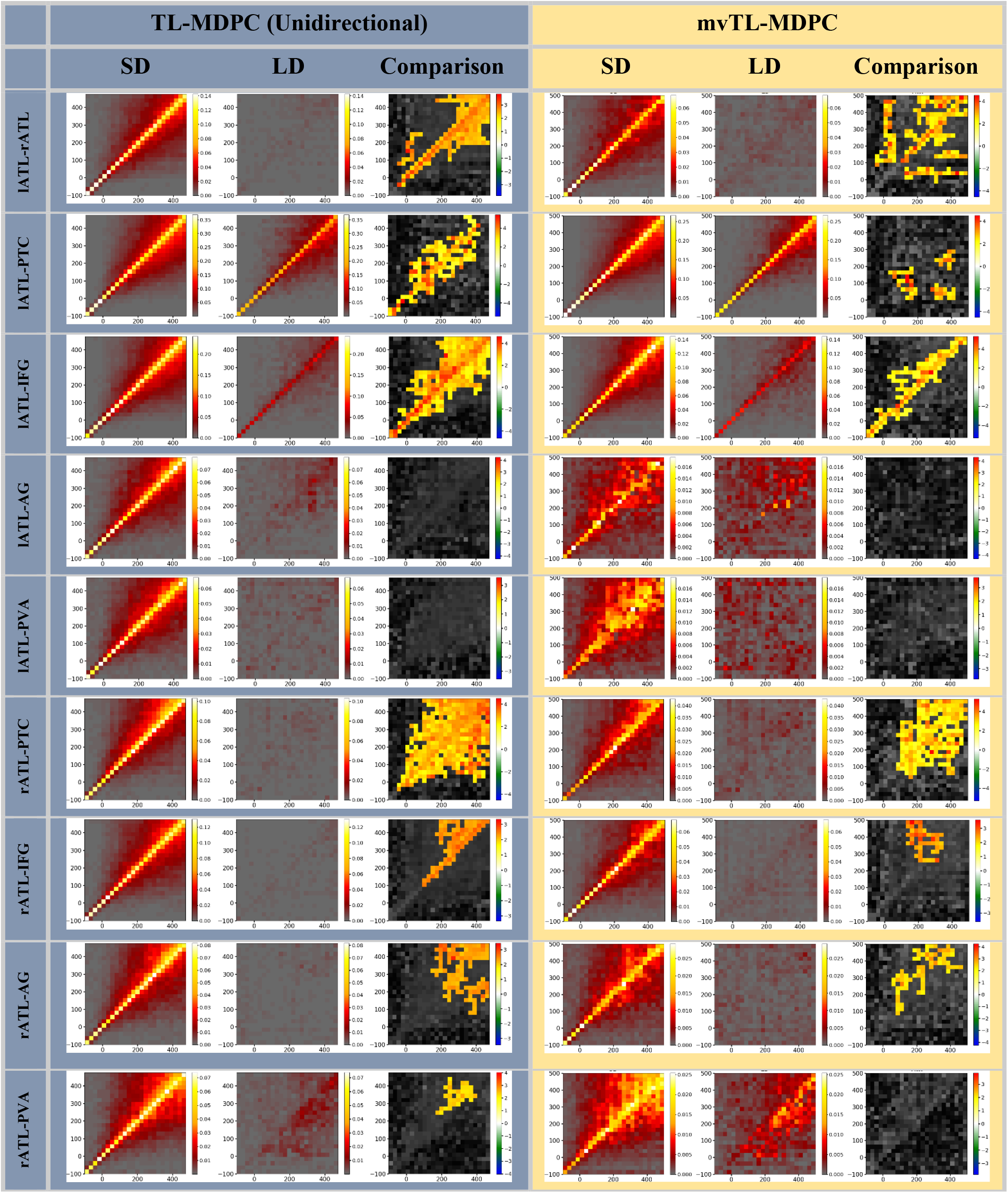
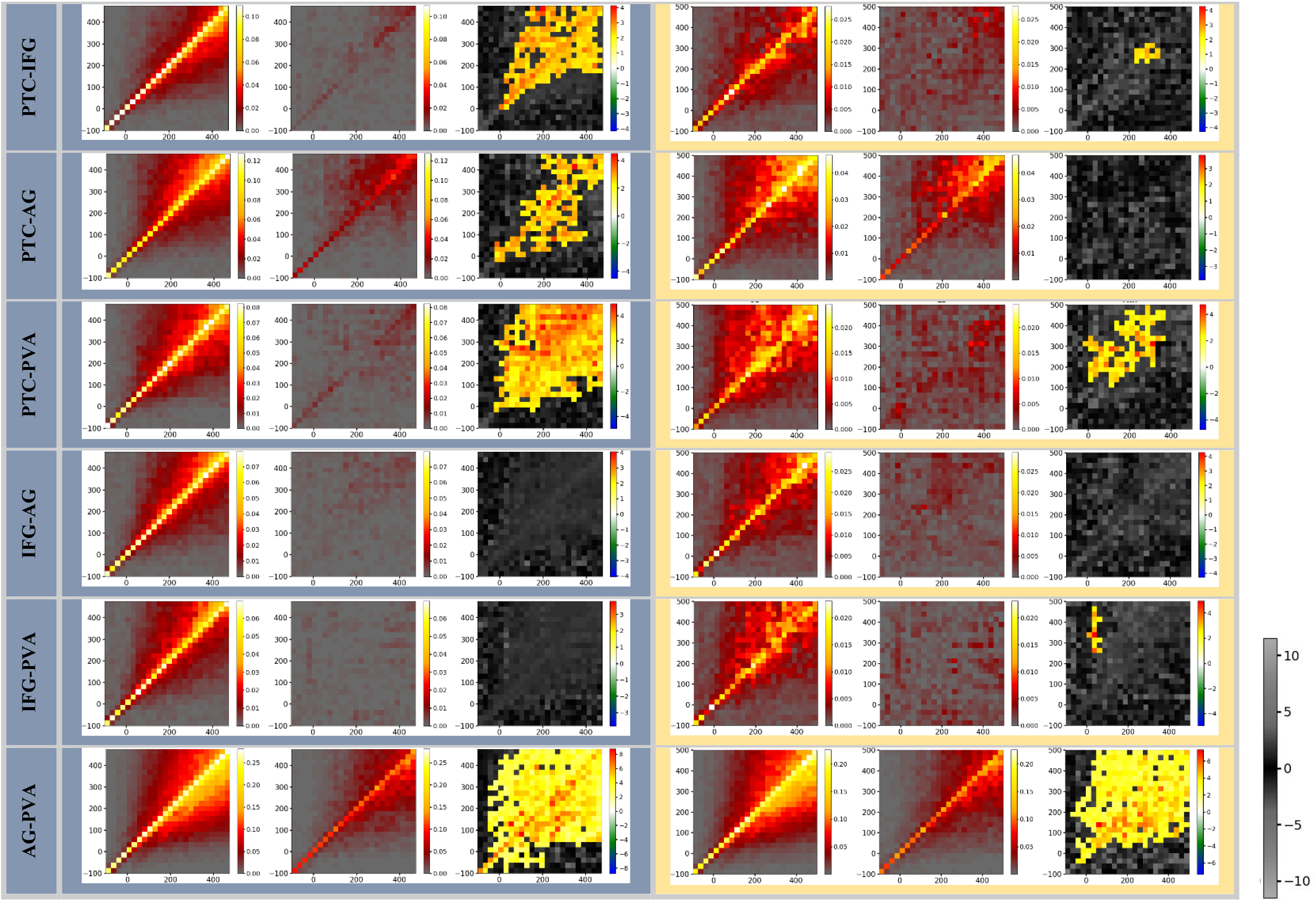
Representation of all TTMs for semantic decision (SD) and lexical decision (LD) tasks, and their comparison, using a unidirectional version of TL-MDPC (left column, computed in one direction to match the multivariate method) and mvTL-MDPC (right column). TTMs for SD and LD are averaged across participants, and comparison was performed using cluster-based permutation test with alpha-level=0.05. All significant contrasts show greater connectivity for SD using MD. To be displayed here, the clusters must be larger than 2% of the TTMs size (24*24). The grey-scale colourbar indicates non-significant t-values (this colour bar is the same across all matrices). Note that results are very similar to the bidirectional version, therefore, difference between the methods are not due to the unidirectional or bidirectional computation of the TL-MDPC.

**Figure S3.**
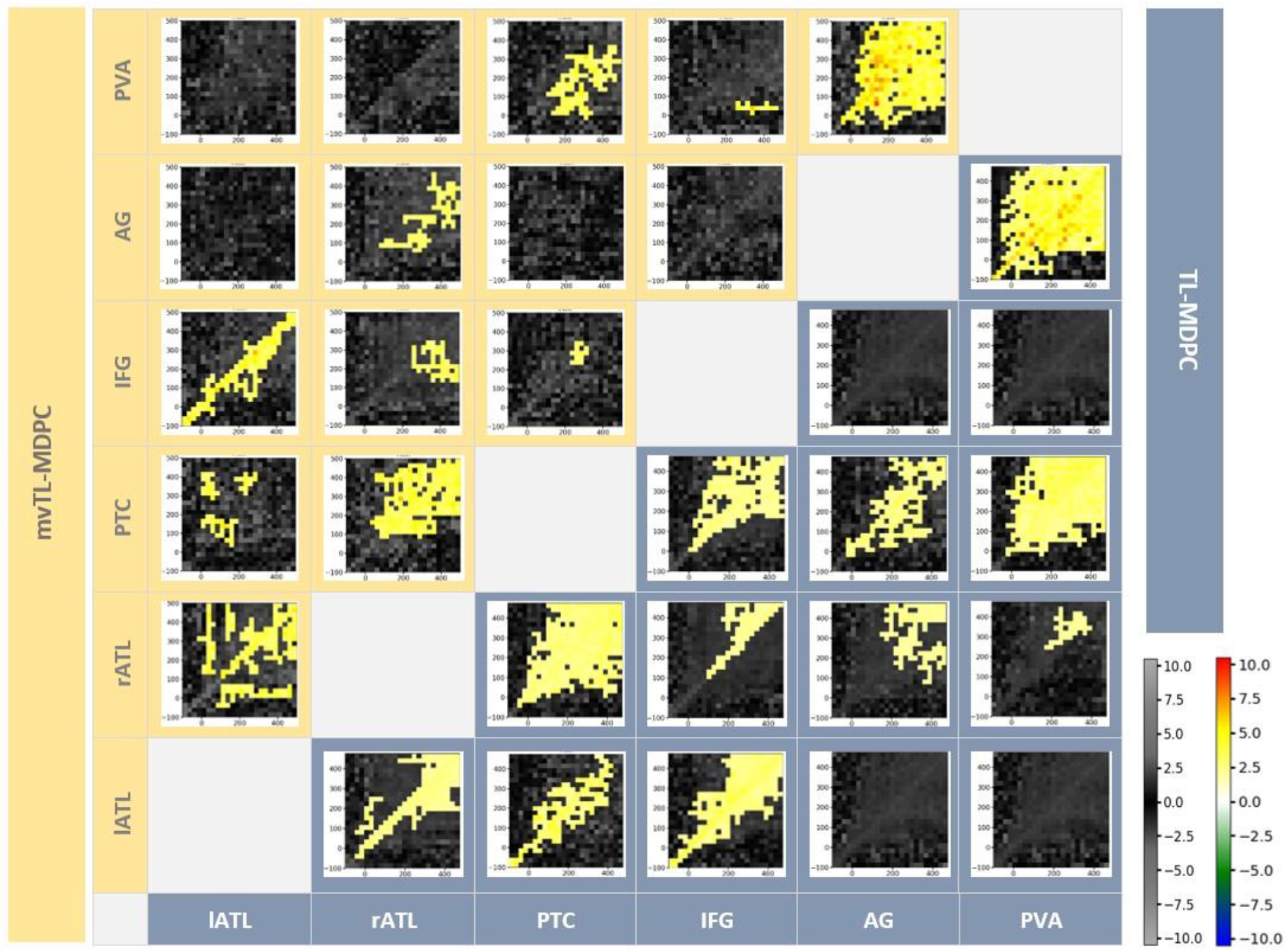
Inter-regional Connectivity Matrix (ICM) for semantic network modulations in the brain with a unidirectional version of the bivariate method – the upper diagonal (yellow shaded area) shows mvTL-MDPC TTMs and the lower diagonal (blue shaded area) shows TL-MDPC TTMs (computed in a single direction to match the multivariate method). Each TTM reflects EVs for a pair of region, ROI X and ROI Y, and across time lags. All modulations obtained by both methods revealed more connectivity for SD than LD. Cluster size was thresholded at 2% of TTMs size (24*24). With both methods, we got rich modulations between semantic control and representation regions. Note that results are very similar to the bidirectional version, therefore differences between the methods are unlikely due to the unidirectional or bidirectional computation of the TL-MDPC.

## Notes

### Competing Interest Statement

The authors have declared no competing interest.

